# Quantifying common and distinct information in single-cell multimodal data with Tilted-CCA

**DOI:** 10.1101/2022.10.07.511320

**Authors:** Kevin Z. Lin, Nancy R. Zhang

**Affiliations:** University of Pennsylvania, Department of Statistics and Data Science

## Abstract

Multimodal single-cell technologies profile multiple modalities for each cell simultaneously and enable a more thorough characterization of cell populations alongside investigations into cross-modality relationships. Existing dimension-reduction methods for multimodal data focus on capturing the “union of information,” producing a lower-dimensional embedding that combines the information across modalities. While these tools are useful, we develop Tilted-CCA to quantify the “intersection and difference of information”, that is, a decomposition of a paired multimodal dataset into common axes of variation that is shared between both modalities and distinct axes of variation that is found only in one modality. Through examples, we show that Tilted-CCA enables meaningful visualization and quantification of the cross-modal information overlap. We also demonstrate the application of Tilted-CCA to two specific types of analyses. First, for single-cell experiments that jointly profile the transcriptome and surface antibody markers, we show how to use Tilted-CCA to design the target antibody panel to best complement the transcriptome. Second, for single-cell multiome data that jointly profiles transcriptome and chromatin accessibility, we show how to use the common embedding given by Tilted-CCA to identify development-informative genes and distinguish between transient versus terminal cell types.

---

High-dimensional multimodal data, where features belonging to two or more modalities are simultaneously profiled, are becoming increasingly widespread across disciplines. In this paper, we focus on *paired* multimodal data arising in the field of single-cell biology, where technological advances have recently enabled simultaneous profiling of multiple types of features, such as RNA expression, protein abundance, and chromatin accessibility all within the same cell and across many cells in parallel^1–4^. This type of data is invaluable because cellular processes operate on multiple molecular modalities, and observation of any single modality offers only a partial view of an inter-connected system^5–9^. In the analysis of multimodal data, a basic question that arises is how to separate and quantify the variations that are shared across modalities and those unique to a particular modality. We address this question and demonstrate how quantifying the shared and unique variations can provide scientific insight. Despite the explicit focus on single-cell genomics, the questions addressed and method developed here are broadly applicable for paired multimodal data in general. For concreteness and clarity, we will refer to the individual data points as “cells.”

Dimension-reduction methods have been useful in analyzing paired multimodal single-cell data by providing a low-dimensional space that typically captures the “union of information” across both modalities. Roughly speaking, these methods seek a low-dimensional embedding of the cells such that a distinct sub-population is distinguishable in the embedding if it is distinguishable in either modality. Methods to estimate such a joint embedding include JIVE^10^, WNN^11^, MOFA+^12^, scAI^13^ and JSNMF^14^. These methods have been useful for identification of nuanced cell types by combining information across both modalities.

In contrast, there is yet no rigorous way to quantify the information that is common to both modalities. Compared to the task performed by the aforementioned methods, this is a fundamentally different task that could be thought of as learning the “intersection of information” between modalities. We define this from a geometric perspective – two sets of cells are separated in the “common” embedding if *both* modalities agree that they are separable. Our matrix factorization method, Tilted-CCA, estimates this embedding by first performing Canonical Correlation Analysis (CCA)^15^ and then optimizing for a suitable decomposition of the canonical score vectors that abides by the aforementioned geometric perspective. We show through examples that Tilted-CCA provides meaningful quantifications of the overlap (or lack thereof) in information between the two modalities at both the cell and the feature level. We note that the “union” and “intersection” embeddings complement one another when analyze multimodal data – while the former provides a complete view of all axes of variation supported by either modality, the latter provides insight on which axes of variation are supported by both modalities. Additionally, Tilted-CCA’s decomposition also quantifies the “distinct” embeddings representing the axes of variation unique to either modality after the intersection has been removed.

We further illustrate the scientific insight enabled by Tilted-CCA through two case studies. In our first case study, we consider the problem of antibody-panel design in paired single cell profiling of RNA expression and surface antibody abundance^16,17^. Such data is becoming standard in immunology research, where modern transcriptome-level cell classifications need to be reconciled with traditional immune cell labels based on surface antibody abundance^18–20^, hopefully giving more accurate labeling of cell identities^21–23^. However, large antibody panels are expensive, especially for studies involving large cohorts. Towards this end, we demonstrate that by quantifying the common and distinct information between the RNA and protein modalities, Tilted-CCA helps in the design of small antibody panels that most effectively separate immune cell types when paired with transcriptomic data.

In our second case study, we investigate the coordination between chromatin accessibility and gene expression during tissue development, and show how Tilted-CCA gives rise to natural metrics for distinguishing between transient versus terminal state cells as well as development-associated genes^24–27^. Many pseudotime-estimation methods have been developed to answer such questions, but they make use of gene expression alone^28–34^ or chromatin accessibility alone^35^, and typically require an estimate of the developmental trajectory as input. In contrast, we show that Tilted-CCA’s common embedding, applied to single-cell transcriptome and chromatin accessibility data, provides an alternate approach to address these questions.

## Results

### Overview of Tilted-CCA, and the intersection of information

We introduce Tilted-CCA through an example of a single-cell CITE-seq data set, where 30,672 human bone marrow cells are simultaneously profiled along the transcriptome (RNA modality) and a panel of 25 surface antibody markers (protein modality)^18^. The modality-specific UMAPs demonstrate that while the major immune subtypes such as myeloid, B-, and T-cells are separated in both modalities, the protein modality better separates the T-cell subtypes such as CD4+ and CD8+ T-cells (Fig. 1a). This is expected since the 25 protein antibodies are chosen to target many T-subtype markers^11^. We use this multimodal dataset to exemplify matrix factorization methods, which strive to factorize the data matrices *X*^(1)^ and *X*^(2)^ of both modalities into the product of modality-specific loading matrices (*L*^(1)^ or *L*^(2)^) and score matrices that decompose into a common embedding *C* and a modality-specific distinct embedding (*D*^(1)^ or *D*^(2)^) (Fig. 1b). However, depending on which axes of variation are represented in the constructions of *C* and *D*, their mathematical properties as well as biological interpretations can vary dramatically.

**Figure 1.**
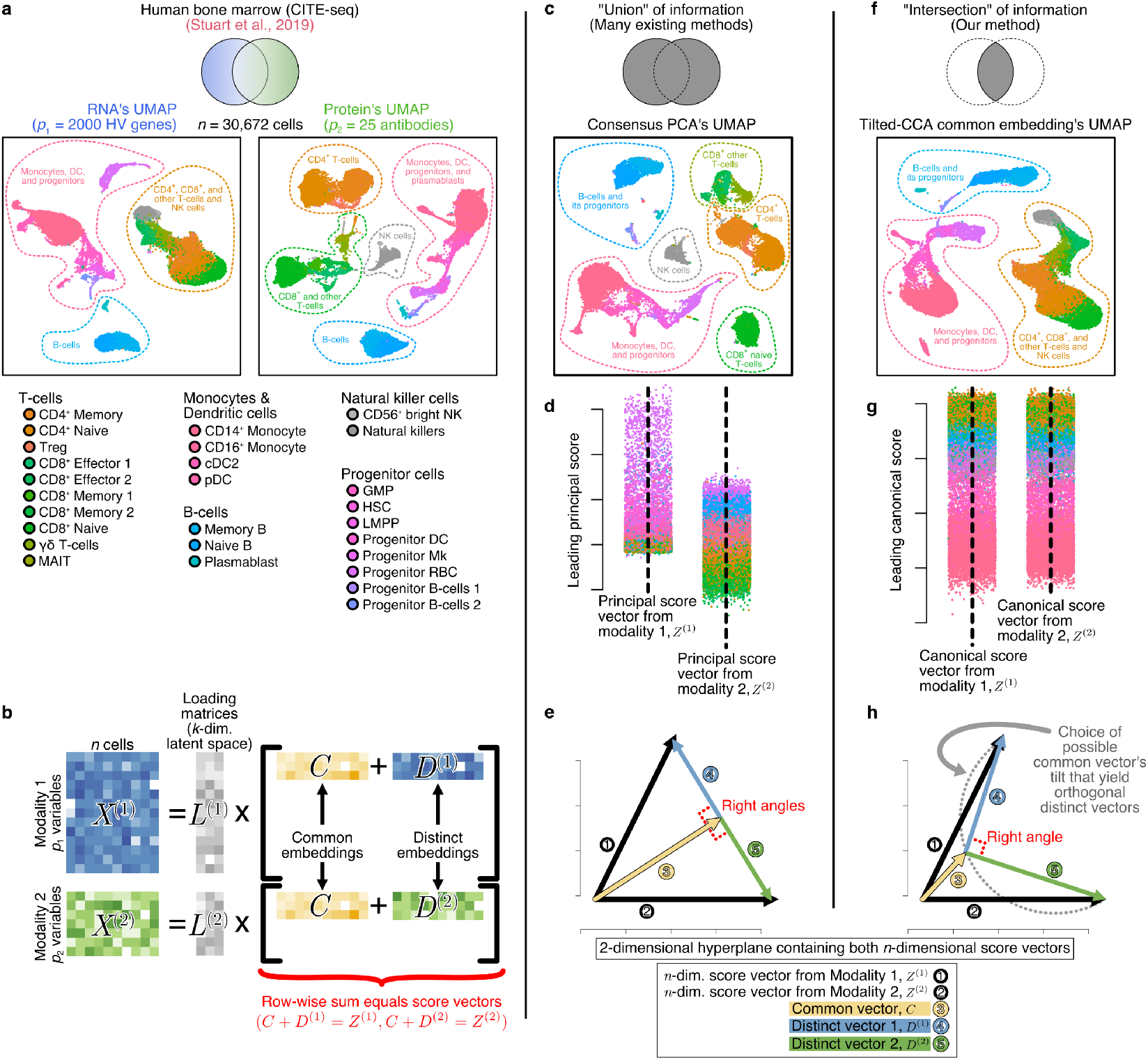
Embedding methods for multimodal data that learn the “intersection” of information differ from those that learn the “union”. **a**, Summary of the bone marrow CITE-seq dataset, showing either the UMAP of the RNA or protein modality, where cells are colored by the annotated cell-types. The coloring scheme persists in **(c,d,f,g)**. **b**, Schematic of a matrix factorization of single-cell paired multimoic data, where 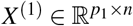 denotes the *n* cells measured on *p*_1_ features and 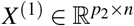 denotes the same *n* cells measured on *p*_2_ features, already preprocessed to be low-rank. Here, *Z*^(1)^ and *Z*^(2)^ denote matrices of score vectors, where the specific definition of these score vectors depends on the context. **c**, UMAP of the global embedding between the RNA and protein modalities estimated by Consensus PCA, showing the “union of information.” **d**, The pair of leading principal score vectors, one from each modality. **e**, Mathematical illustration of the decomposition of *Z*^(1)^ and *Z*^(2)^ (the two sets of principal score vectors) to achieve a decomposition as shown in (**b**) as used by methods like JIVE. **f**, UMAP of the common embedding *C* between the RNA and protein modality estimated by Tilted-CCA, showing the “intersection of information.” **g**, The pair of leading canonical score vectors, one from each modality. **h**, Mathematical illustration of the decomposition of *Z*^(1)^ and *Z*^(2)^ (the two sets of canonical score vectors) used by Tilted-CCA to achieve a decomposition as shown in (**b**).

Broadly speaking, existing methods define C in Figure 1b to capture the “union” of information^10^. A prototypical method is Consensus Principal Component Analysis (PCA)^36–38^, which finds a low-dimensional embedding to best approximate both data matrices *X*^(1)^ and *X*^(2)^ by combining the leading PCA axes of variation from each modality. For this CITE-seq dataset, Consensus PCA simultaneously separates all of the progenitor cell-types, as well as the CD4+, CD8+ and NK T-cells from each other (Fig. 1c). To explain how Consensus PCA yields a decomposition in the form of Figure 1b, consider the leading principal component from each modality, Z^(1)^and Z^(2)^, each of which differentiates a different set of cell types (Fig. 1d). The common embedding C is the linear subspace that best retains the cell-type separation patterns in both principal components, and *D*^(1)^ and *D*^(2)^ are the residuals orthogonal to C which are interpreted as the modality-specific distinct embeddings (Fig. 1e). Other linear methods such as JIVE^10^, MOFA+^12^, scAI^13^ and JSNMF^14^ and non-linear methods such as WNN^11^ have similar qualities which learn the leading axes of variation from either modality (Extended Data Fig. 1, see Supplementary Information for more details).

In contrast to the aforementioned methods, Tilted-CCA finds a low-dimensional embedding that quantifies axes of variation that are supported by both modalities in an unsupervised fashion, i.e., the “intersection” of information. For example, in this CITE-seq data, this embedding separates the myeloid, B-, and T-cells from one another, since this separation is supported by both modalities, but leaves the signal that separates the CD4+ and CD8+ T-cell subtypes to the distinct embedding, since their separation is unique to the protein modality (Fig. 1f). As the name suggests, Tilted-CCA builds on CCA, which finds linear transformations for each modality that have the highest cross-modal correlation. Here, the leading pair of canonical vectors in CCA exemplifies the shared pattern between both modalities that separate the most major cell types (Fig. 1g). However CCA, by itself, only provides two canonical score matrices *Z*^(1)^ and *Z*^(2)^, and does not imply an explicit decomposition of the common and distinct axes of variation in the framework of Figure 1b. Tilted-CCA fills this gap by starting from CCA and then decomposing *Z*^(1)^ and *Z*^(2)^ into a common embedding C and two distinct embeddings *D*^(1)^ and *D*^(2)^ where: 1) C encapsulates the appropriate geometric relations among the cells that are supported by both modalities, and 2) *D*^(1)^ and D^(2)^ are constrained to be orthogonal to each other. The latter orthogonality constraint was first proposed in D-CCA^39^, and it enables us to interpret *D*^(1)^ and *D*^(2)^ as capturing modality-specific variation and ensures that axes of variation in C have magnitudes that are proportional to the canonical correlation. This constraint *D*^(1)^ ⊥ *D*^(2)^ restricts the common vector in each latent dimension of C to lie along a semi-circle defined by the canonical score vectors Z^(1)^ and Z^(2)^ (Fig. 1h). Along this semi-circle, if *C* tilts in the direction of *Z*^(1)^, the common embedding would resemble Modality 1, leading to the interpretation that Modality 2 has more distinct information represented by large magnitudes in *D*^(2)^, and vice versa (Fig. 2a). Thus, Tilted-CCA searches for the appropriate “tilt” of *C* along this semi-circle that yields the desired geometric relations among the cells. Specifically, Tilted-CCA first computes the nearest-neighbor graph of each modality where each node is a cell (Fig. 2b). Then, a target common manifold is constructed from both graphs in order to encapsulate the cell-type separation patterns common to both modalities – broadly speaking, two cells are separated in this manifold only if both modalities agree that they should be separated. Tilted-CCA optimizes the tilt of the common vector for each latent dimension so that the resulting common embedding’s nearest neighbor graph approximates this target manifold (Fig. 2c). Once the appropriate common embedding C is estimated, we recover the estimated matrix decomposition of both distinct embeddings in the framework of Figure 1b (Methods).

**Figure 2.**
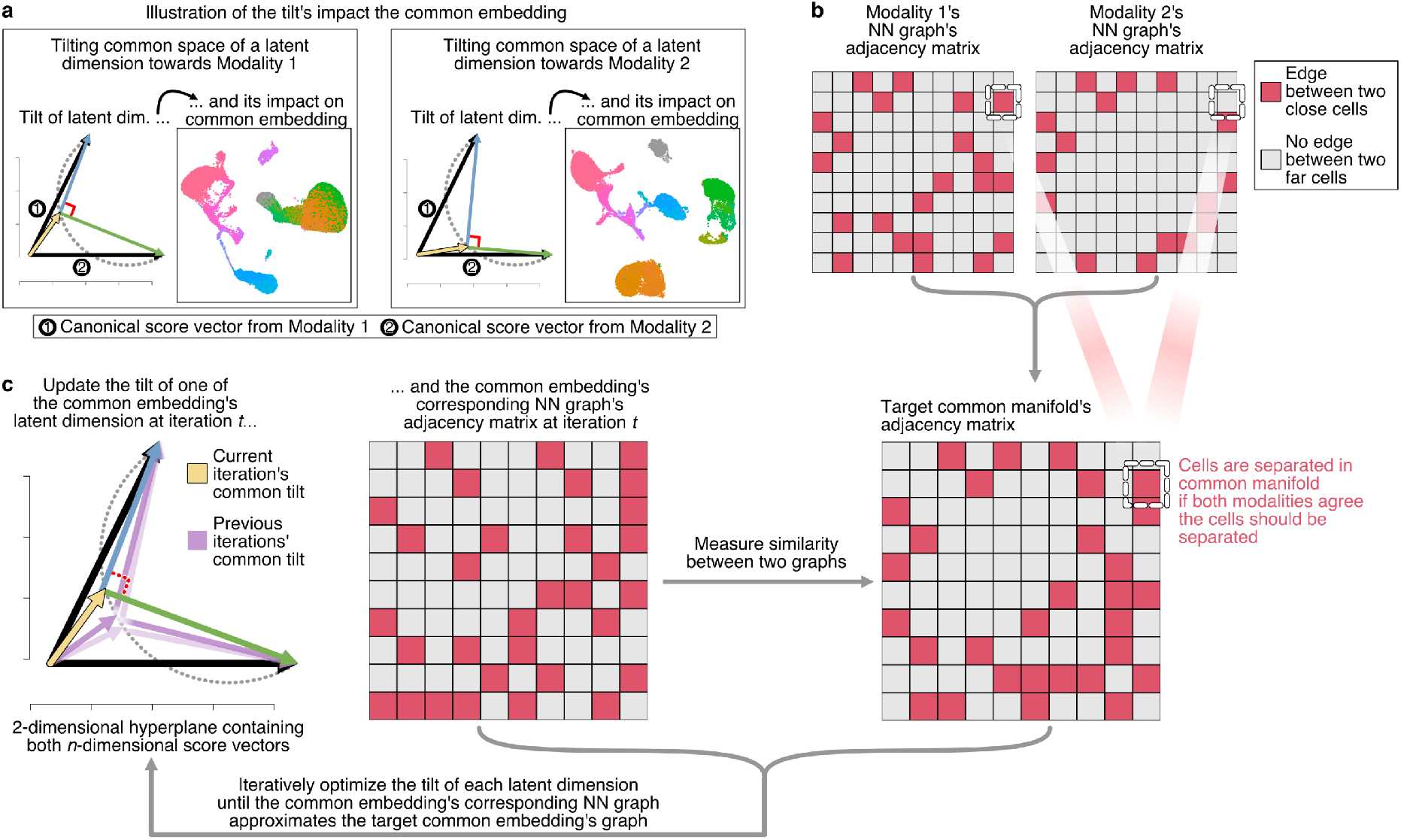
Schematic of Tilted-CCA’s optimization of common embedding’s tilt across each latent dimension. **a**, Schematic illustrating the tilt’s impact on the common embedding using the bone marrow CITE-seq dataset, where the tilt for a latent dimension could either lean towards Modality 1 (meaning Modality 2 has more distinct information in this latent dimension) or vice versa. **b**, Schematic of the nearest-neighbor graphs for each modality. **c**, Flowchart of Tilted-CCA’s optimization procedure. The target common manifold (represented as an adjacency matrix) is computed based both modalities’ respective nearest-neighbor graphs. Then, an iterative optimization procedure is used where the tilt for each latent dimension is updated based on how similar its corresponding common embedding’s nearest-neighbor graph is similar to the target common manifold.

### Titled-CCA quantifies the overlapping versus distinct information each modality contributes towards the separation of cell types

As proof of concept, we start with a pervasive question asked for multimodal single-cell data: which cell types are distinguishable by both modalities, and which are distinguishable by only one modality? We use the cell-type annotations provided by the authors to investigate this. For example, the UMAPs of the RNA and protein modalities for the aforementioned CITE-seq bone marrow dataset suggest that the protein modality better differentiates the CD4+ and CD8+ T-cell subtypes than the RNA modality. (Fig. 1a). Can we rigorously quantify the degree to which this is true, and survey the amount of overlapping (i.e., shared) versus distinct information each modality contributes towards the separation of predefined cell types in the data?

Prior to addressing these questions quantitatively, we qualitatively explore the UMAPs of *D*^(1)^ and *D*^(2)^, the distinct components derived from the orthogonal remainder terms after removal of the common embedding C. For the bone marrow CITE-seq data, we see that CD4+ and CD8+ T-cells are mixed in the RNA’s distinct component *D*^(1)^, while the progenitor cell-types are well separated (Fig. 3a). On the other hand, the progenitor cell-types mixed in the protein’s distinct component *D*^(2)^, while the CD4+ and CD8+ T-cells are well separated. These observations match our initial intuitions gained from a side-by-side comparison of each modality’s UMAP in Figure 1a alone. The major cell types such as myeloid cells, B- and T-cells are also distinguishable in both distinct UMAPs, indicating that there are remaining axes of variation in both modalities that separate these cell types, but are orthogonal to each other.

**Figure 3.**
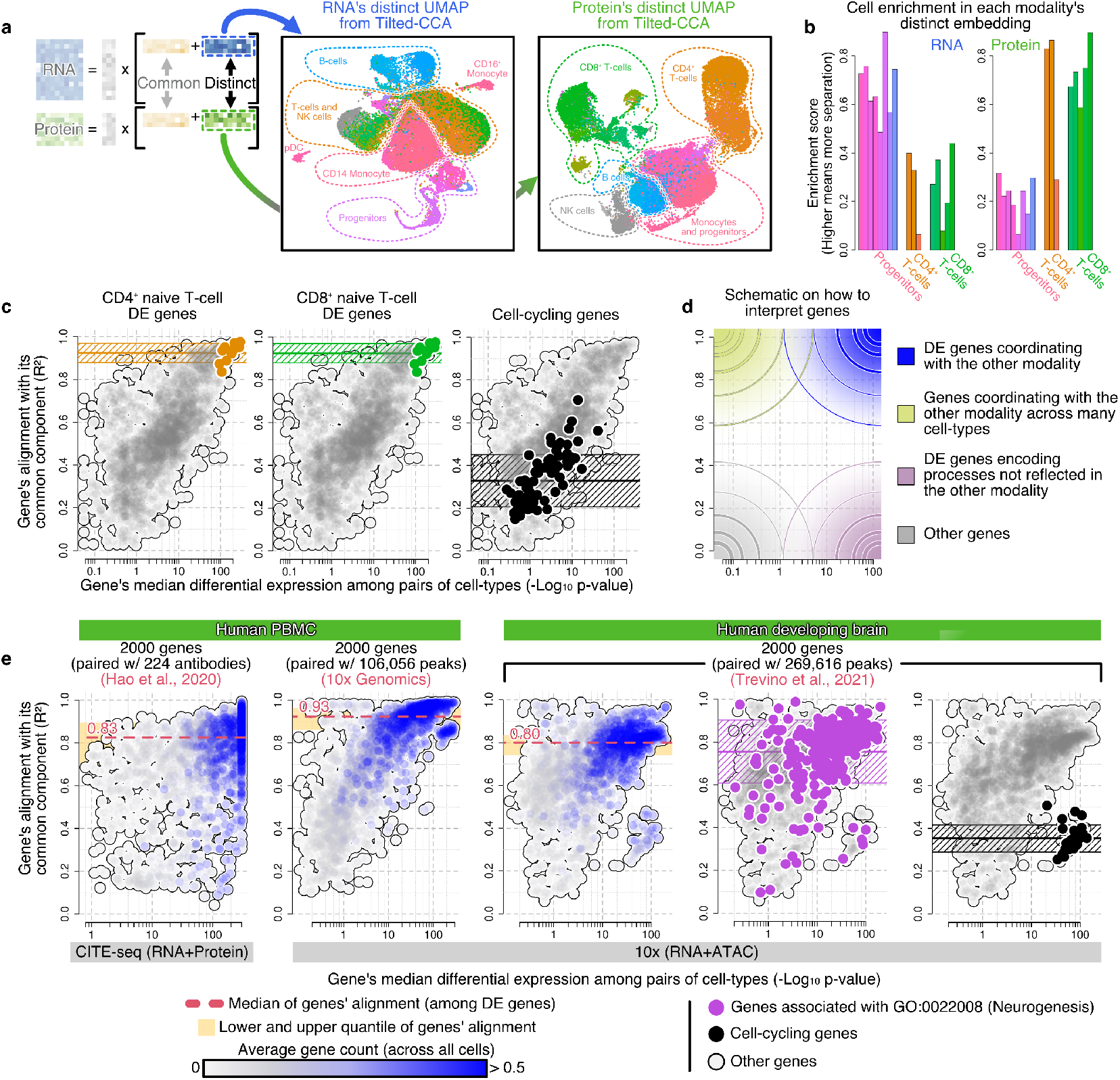
Tilted-CCA quantifies shared/distinct information on both the cell- and feature-level, enabling within-dataset and between-dataset comparisons. **a**, Schematic of Tilted-CCA’s matrix factorization and UMAP of each modality’s distinct embedding in the bone marrow CITE-seq dataset. **b**, Enrichment score of various cell-subtypes among progenitors, CD4+ T-cells and CD8+ T-cells based on either the RNA’s or protein’s distinct embedding. **c**. Alignment-differentiability plot of the 2000 genes for the bone marrow CITE-seq dataset, marking either marker genes for CD4+ naive T-cells, CD8+ naive T-cells or cells associated with cell-cycling. The median and quantiles of the alignment are shown along the y-axis. **d**, Schematic of how different regions of the alignment-differentiability plot are informative of different features’ qualities. **e**, Alignment-differentiability plots of the 2000 genes for various datasets measuring RNA+protein or RNA+ATAC modalities, where genes are colored based on their average count and the median alignment among DE genes for each dataset is marked by a red dashed line. Additionally, the genes associated with neurogenesis and cell-cycling are marked in purple and black respectively, with the median and quantiles among these sets of genes shown along the y-axis.

Without relying on UMAPs, the amount of distinct information contributed by each modality towards the separation of a cell type can be formally quantified using *enrichment scores*: first, compute two nearest-neighbor graphs, one for each of *D*^(1)^ and *D*^(2)^. Then, define the enrichment score of a cell-type for a particular distinct embedding by selecting all cells of said type and computing the proportion of their neighbors that share their type, normalized by the baseline proportion (Methods). The enrichment scores quantify which modality contains more distinct information towards the separation of each cell type. We see that for the bone marrow CITE-seq data, the progenitors are roughly 3 times more separated in RNA’s distinct space as compared to in protein’s distinct space, while the CD4+ and CD8+ T-cells are roughly 2.5 times more enriched in protein’s distinct space as compared to in RNA’s distinct space (Fig. 3b). While the fact that the protein modality contains more information that separates the T subtypes is visible in Figure 1a, the fact that the RNA modality contains more distinct information separating the progenitor cells is not immediately discernible in Figure 1a. This makes biological sense since progenitor cell populations involve transcriptome-level changes not expected to be captured by the 25 antibody markers. We also define the enrichment score for the common embedding, and quantify that the overlapping information between both modalities clearly define the B-cells (Extended Data Fig. 2a,b).

### Tilted-CCA quantifies the degree of cross-modality alignment of features

Now consider the features in each modality, such as the genes in the RNA modality and the surface antibody markers in the protein modality, for the bone marrow CITE-seq data. How much of each feature’s variation lie in the common space versus its modality’s distinct space? Compared to standard PCA analysis, this is analogous to asking which features are most aligned with each principal component, which could be address via the principal loadings. Here we propose an intuitively similar metric for Tilted-CCA: for each feature, we quantify its *alignment score*, defined as the *R*^2^ of regressing its observed expression onto its common component. A higher *R*^2^ implies that the feature’s variation lies mostly in the shared common space.

As proof of principle, which genes in the RNA modality do we expect to align the most with the common space in the bone marrow CITE-seq data? We hypothesize that cell-type marker genes should be at the top of this list, since these genes’ expressions should correlate with the expression of the panel’s antibody markers. To investigate this, we plot the common-alignment *R*^2^ of each gene against its differentiability, a meta-statistic that summarizes its differential expression across all pairs of cell types. A higher differentiability implies that the gene is a strong marker for certain cell type(s). This calculation requires an a priori partitioning of the cells, which can be derived from either an unsupervised clustering or a manual or reference-based assignment of cell types^18^. We see that highly-differentiable genes, such as the markers for CD4+ and CD8+ T-cells, are indeed highly aligned with the protein modality (Fig. 3c). In contrast, genes belonging to house-keeping processes which do not differentiate cell types, such as cell-cycle, have low alignment with common space (Fig. 3c, Extended Data Fig. 2c). This analysis shows that the feature-level alignment to the common space is in accordance with our expectations. See Supplementary Information for analogous analyses, but focusing on the antibodies.

The alignment-differentiability plots also provide a bird’s-eye view on the amount of overlapping information between modalities when comparing across different technologies and biological systems (Fig. 3d). Genes in the upper-right quadrant (blue) of the alignment-differentiability plot are cell-type markers that coordinate with the other modality, while genes in the lower right quadrant (purple) are cell-type markers that complement the other modality. As an example, consider the alignment-differentiability plots for two multiomic experiments profiling PBMC: a CITE-seq experiment pairing full-transcriptome RNA-seq with 224 antibody markers, and a 10x Multiome experiment pairing full-transcriptome RNA-seq with ATAC^40^, which measures chromatin accessibility across the genome (Fig. 3e). The median alignment of differentially-expressed genes (i.e., genes with a differentiability score of more than 10) with the protein modality is remarkably lower than their alignment with the ATAC modality (0.83 and 0.93 respectively). This demonstrates that for PBMC, the transcriptome provides additional cell-type separation patterns not present among the 224 antibody markers, but the transcriptome and chromatin accessibility predominately capture the same axes of variation. Next, compare two different tissues, PBMC and developing brain, both sequenced using 10x Multiome. The cross-modality alignment is much higher in PBMC as compared to developing brain (0.93 and 0.8 respectively). Although some of this difference is due to slight differences in ATAC sequencing coverage between the datasets (Extended Data Fig. 2d), we hypothesize that the main difference is biological – the developing brain contains mostly differentiating cell populations, where RNA expression is less in sync with chromatin-level differences as compared to PBMC, which is a terminally-differentiated population. This is supported by examining genes relevant to neurogenesis, many of which have low alignment with the common space. This observation suggests that the degree of feature alignment with Tilted-CCA’s common embedding may contain developmental information, which we examine in detail a later section through the construction of synchrony scores.

It is also worth noting, from Figure 3e, that cell-cycle genes in general have high differentiability and low common space alignment. The fact that the cell-cycle signal is unique to the RNA modality and not shared with ATAC, is a recurring theme in paired RNA and ATAC multiome sequencing of developing tissues (Extended Data Fig. 2e). This makes biological sense as cell-cycle is a transient process and not a permanent aspect of cell identity, and we expect only the more permanent changes related to cellular identity to be encoded at the chromatin level.

### Designing minimally-sized antibody panels for CITE-seq data with Tilted-CCA

Multiomic sequencing technologies for joint assay of RNA expression and surface protein abundancy, such as CITE-seq^16^ or Abseq^17^, require practitioners to select the antibody panel (see Extended Data Fig. 3 for additional examples). Large antibody panels are expensive, making them impractical for large cohort studies. Hence, we desire designing a small panel of antibodies that provide the most distinct information, complementary to RNA, for the purpose of separating cell types. We hypothesize that Tilted-CCA is suitable for achieving this, since quantifying the intersection and difference between two modalities can aid in selecting the features that best contribute to their union.

For illustration, we consider an Abseq dataset of 461 genes and 97 surface antibodies of cells from human bone marrow^20^ (Fig. 4a). Figure 4b previews the panel of 10 antibodies selected by Tilted-CCA (see also Extended Data Fig. 4a). Although the 10 antibodies do not give a good separation between cell types in isolation, the Consensus PCA combining these antibodies with the RNA modality yield cell-type separations almost as good as Consensus PCA applied to the full data set containing 97 antibodies (see also Fig. 4c, Extended Data Fig. 4b). At a high level, our antibody-panel design strategy greedily finds a set of antibodies whose protein’s distinct component are differentially expressed across cell types (as defined in the previous section) and are as uncorrelated as possible to each other and to the common embedding (see also Extended Data Fig. 4c). As a diagnostic, we plot the correlation network among the 97 antibodies, where edges connect pairs of antibodies with highly correlated distinct components (Fig. 4d). Here, the node color and size reflect differentiability and alignment with common space respectively. The 10 selected antibodies are spread out across this network, demonstrating their uncorrelated nature, while having highly-differentiable distinct components that are lowly aligned with the common subspace. In contrast, conventional immune markers such as CD11b+, CD14+ and CD16+ are not selected since their protein expression are already aligned with RNA.

**Figure 4.**
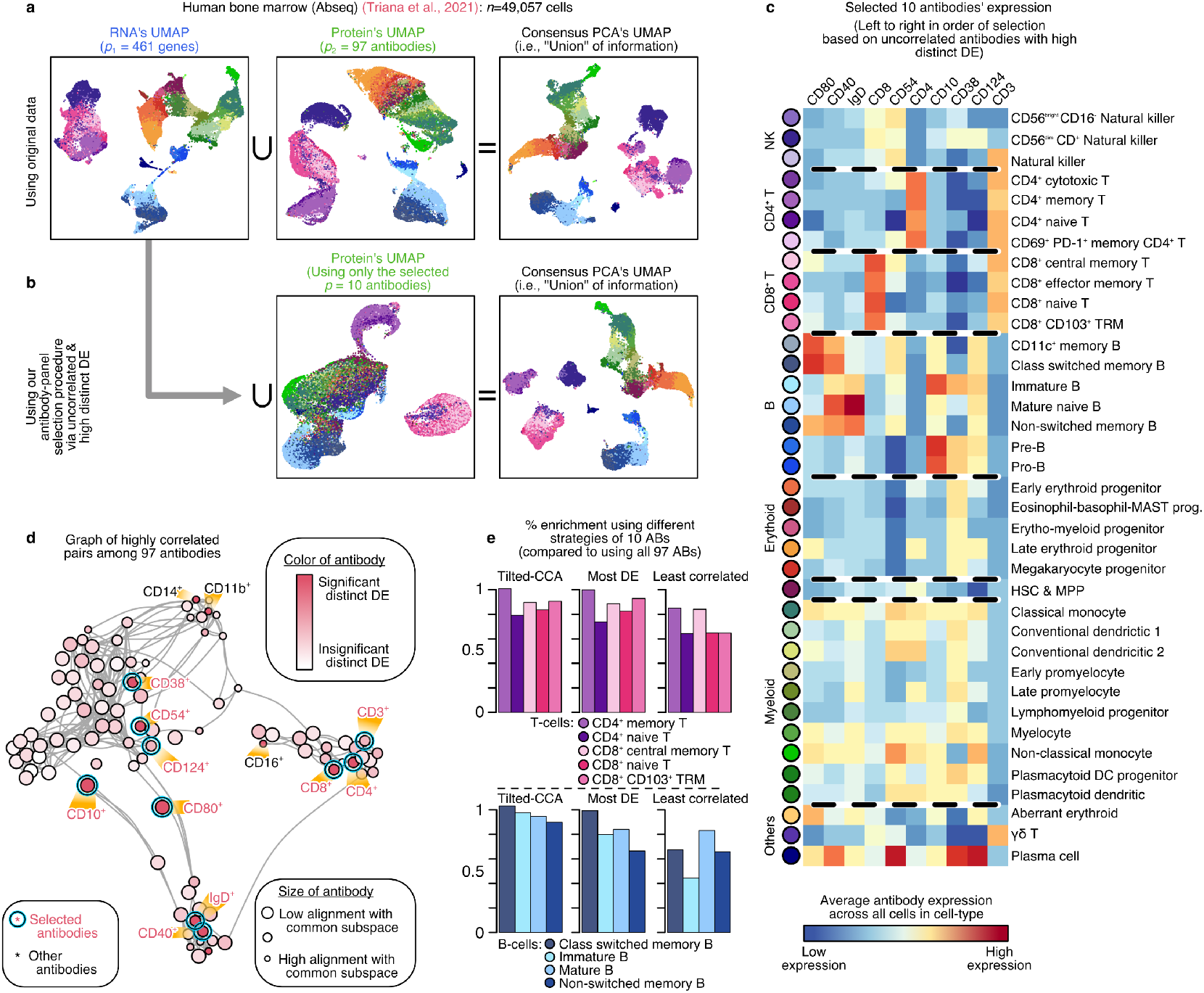
Tilted-CCA enables targeted antibody panel design for RNA+protein multiome assays. **a**, UMAP of the RNA and protein modalities, as well as the Consensus PCA capturing the union of information. **b**, UMAP of the protein modality restricted to the 10 selected antibodies using our procedure showing poor separation among annotated cell-types, and the resulting Consensus PCA when combined with the RNA modality showing many of axes of variations compared to (**a**). **c**, Heatmap of the 10 selected antibodies across the different cell types, where antibodies are ordered by order of selection from left to right. **d**, Correlation graph among the 97 antibodies, where the size of a node denotes the antibody’s alignment with Tilted-CCA’s common subspace and the color denotes the significance of the antibody’s distinct component. All 10 selected antibodies are marked, as well as other conventional immune markers that were not selected by our procedure. **e**, Percent enrichment of different cell types (either various CD4+/CD8+ T-cells or B-cells) of the Consensus PCA obtained by combining the RNA modality with different panels of 10 antibodies (obtained either using our Tilted-CCA, the most differentially-expressed antibodies, or the least correlated with the transcriptome), when compared to the enrichment of the Consensus PCA obtained by combining the RNA modality with all 97 antibodies. A higher enrichment percent denotes less information is lost when using 10 antibodies.

As comparison, we also consider two other strategies for antibody-panel design: selecting the 10 antibodies that are most differentially-expressed, or the 10 antibodies that are least correlated with the RNA modality. These methods yield Consensus PCA embeddings that have poorer separation among cell types, as quantified by using the aforementioned enrichment scores (Fig. 4e, Extended Data Fig. 4d,e). This is particularly evident among CD4+/CD8+ T-cells and B-cells. The former strategy suffers since highly-differentiable antibodies might have expression patterns already prevalent in the gene expression, and thus provides redundant information when paired with the RNA modality. The latter strategy suffers since antibodies not correlated with gene expressions do not necessarily provide high cell-type separation.

### Tilted-CCA reveals transient cell states and development-informative genes in developing cell populations

Joint profiling of gene expression and chromatin accessibility at the single-cell level enables the study of the coordination between chromatin remodeling and transcriptome reprogramming during cellular differentiation^26,41^. Towards this end, we explore the use of the common and distinct embeddings of Tilted-CCA to answer two specific questions: first, can we identify which cells are in a transient or terminal cell-state? Second, can we identify genes that are associated with development, and characterize the temporal coordination between the chromatin activity and RNA expression of these genes? Many pseudotime-estimation methods, based on only RNA or ATAC alone, have been developed to address these questions, and these methods typically start by estimating the underlying cell trajectory and/or RNA velocity fields^28–30,32,35,42,43^. Trajectory estimation requires cell differentiation signals to be strong in the data, and reliable trajectories are often difficult to recover with confidence. Complementing existing pipelines, we take an alternative approach to address the above two questions that does not depend on the estimation of cell trajectory or RNA velocity field.

We start with the premise that development is characterized by a coordinated change between the transcriptome and chromatin accessibility. Hence, we posit that the geometry of cells in the common embedding between RNA and ATAC captures the differentation trajectory, and that large deviations from the common embedding capture the asynchrony between RNA-level and chromatin-level signals. The coordinated yet asynchronous change between chromatin remodeling and gene transcription has been characterized by numerous recent studies on transcription priming^26,44–48^. To measure the synchrony of the RNA and ATAC modalities along a latent developmental trajectory (without knowledge of that trajectory), we design a linear regression to predict the RNA common-space component of each gene using the ATAC modality, where large absolute residuals are indicative of a lack of synchrony between that gene’s RNA and ATAC (Methods). The absolute residuals of this prediction are plotted against developmental pseudotime for specific genes relevant to cortical neurogenesis based on a 10x Multiome dataset of developing human brain^40^ (Fig. 5a). Note that the pseudotime is used only for comparison, and not for computing the regression. We see that, indeed, for the neurogenesis-related genes, the residuals are small in terminal cells (i.e., pseudotime close to 1) and large for cells in transition. Encouraged, we design a cell-wise synchrony score, which summarizes the magnitude of these residuals across all genes for each cell, to distinguish between cells in terminal versus transient state without the reliance on trajectory reconstruction and pseudotime estimation (Methods).

**Figure 5.**
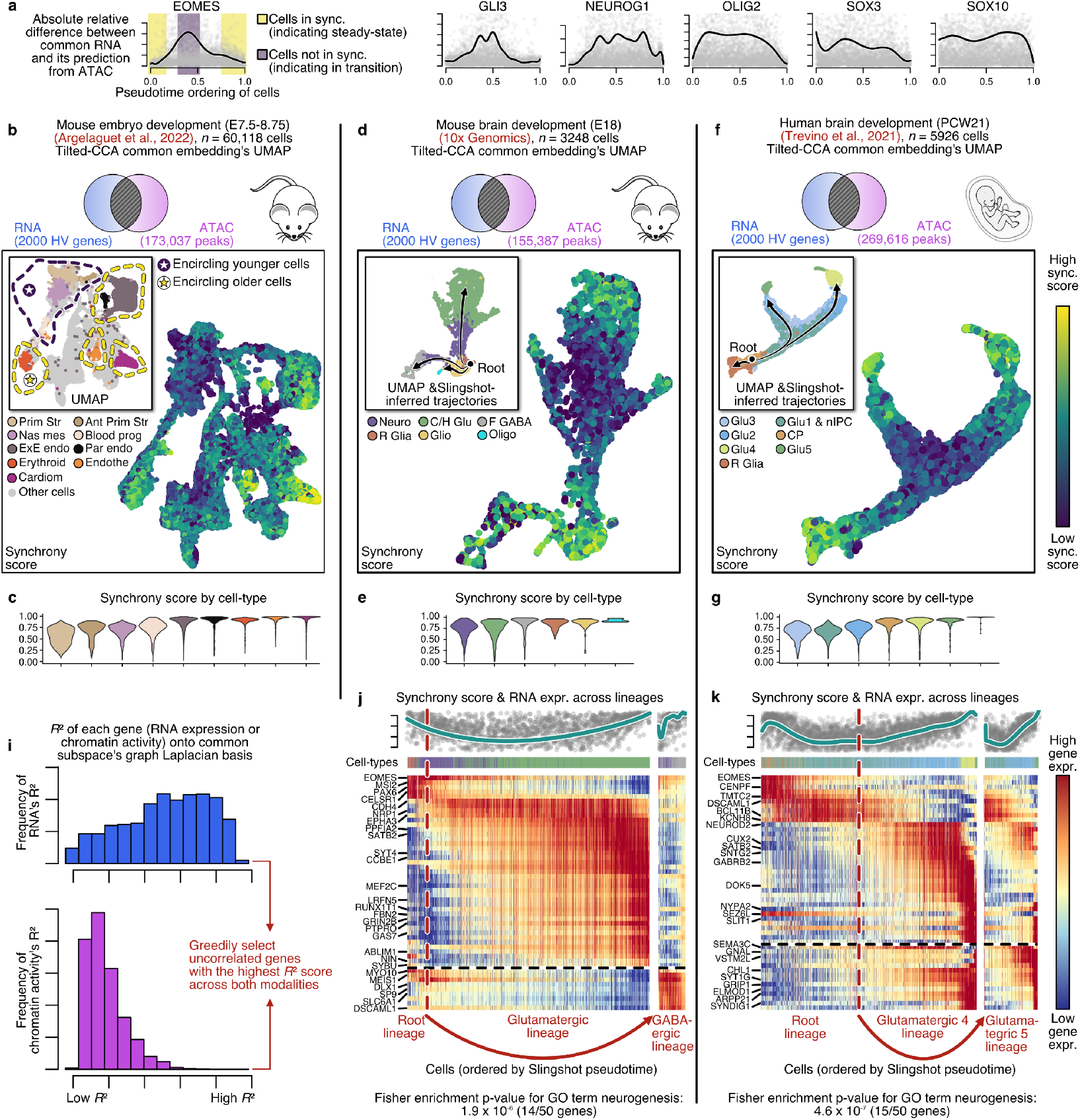
Tilted-CCA infers cell’s developmental status and development-associated genes based on the common embedding between the RNA and ATAC modality. **a**, Asychrony between the ATAC and RNA measured by the residual of predicting the common RNA with the ATAC modality, plotted against Slingshot’s pseudotime ordering of the cells in the Glutamatergic 4 lineage of the human brain development dataset. **b**, UMAP of a mouse embryo dataset where cells are colored based on annotated cell-types (where regions of young cells are circled in purple, and regions of older cells are cicled in yellow) or Tilted-CCA’s synchrony scores. **d,f**, UMAPs of a mouse brain development or human brain development system shown with the Slingshot trajectories where cells are colored based on annotated cell-types or Tilted-CCA’s synchrony scores. **c,e,g**, corresponding violin plots of the synchrony scores across cell types for the respective datasets, illustrating lower synchrony scores for cell-states in transition. **i**, Schematic illustrating the selection procedure for development-associated genes. **j,k**, Heatmaps of selected genes and cells in the glutamatergic lineage or the GABAergic lineage in the mouse brain development system, or selected genes and cells in the Glutamatergic 4 or 5 lineage in the human brain development system. The genes in (**j**) and (**k**) are ordered to visualize the developmental cascade, and the exact Fisher enrichment of the enrichment of the selected 50 genes for the GO term “neurogenesis” is marked. The cells are ordered based on the pseudotimes estimated by Slingshot for visual clarity, and is not required to construct the synchrony scores or the select the development-associated genes.

We apply the synchrony score to three 10x Multiome datasets of developing tissues. First, consider the developing mouse embryo^49^ at embryonic day 7.5 to 8.75, equipped with cell-type labels delineating the youngest cell types such as primitive streak and blood progenitors as well as the terminal cell types such as cardiomyocytes and erythroids. We see that the synchrony scores are indeed low for the former group while high for the latter group (Fig. 5b,c). Next, consider the developing mouse brain at embryonic day 18, where cell-type labels were transferred from an independent RNA reference^50^ using SAVERCAT^51^. Here, cell lineages originate from the radial glia and differentiate into oligodendrocytes, cortical/hippocampal glutamategeric and forebrain GABAergic cell types. We see that both the radial glia and the cells at terminal fates (GABAergic and glutamatergic neurons as well as oligodendrocytes) have high synchrony scores (Fig. 5d,e). In contrast, the neuroblast cells and cells in the earlier stages of glutamatergic differentiation have lower synchrony score. Lastly, consider the developing human brain^40^ at post-conception week 21. Here, the development originates from the cycling progenitors and differentiate into either the radial glia or different types of cortical gluamatergic neurons. The synchrony scores are high for both the cycling progenitors as well as the radial glia and the mature terminal cell types (Fig. 5f,g), and low for the cells in transition. Thus, for all three datasets from developing tissues, the Tilted-CCA synchrony score reliably distinguished between cells in a transient or terminal cell state. Importantly, the synchrony score uses RNA-ATAC relationships not used by existing methods and does not require the a priori estimation of cell trajectory or RNA velocity, hence providing orthogonal information that complements existing methods.

We move on to the second question – without a priori knowledge of the developmental trajectory, can we identify the development-associated genes and characterize the relationship between their ATAC-derived chromatin activity and their RNA expression? Here, we define the chromatin activity as the total read coverage in peak regions from the ATAC modality that are ±500 base pairs from the gene’s transcription start sites. We apply Tilted-CCA to these RNA and chromatin activity modalities, yielding embeddings similar to those in Fig. 5d,f (Extended Data Fig. 5c,e and Extended Data Fig. 6a). Based on the premise that development-associated processes should be captured in Tilted-CCA’s common embedding, we measure how well each gene’s expression and chromatin activity conform to the geometry of the common embedding’s nearest-neighbor graph. To get a representative list of gene markers for all stages and branches of development, we greedily select genes that highly conform to the common embedding’s geometry in both modalities and are uncorrelated with each other (Fig. 5i, Methods). The 50 genes identified in this way for both the mouse and human developing brain systems are highly enriched for neurogenesis and display varied expression profiles across pseudotime and lineages (Fig. 5j,k, and Extended Data Fig. 6b). Importantly, the ordering of cells in the shown heatmaps are chosen based on Slingshot’s pseudotime, and is used only for visualization but not selection. We also show the synchrony score of each cell against its ordered pseudotime, indicating that, as expected, the synchrony is indeed low for the transitioning cells. These findings demonstrate that Tilted-CCA’s common embedding provides an alternative method of selecting development-associated genes that do not require the prior estimation of the developmental trajectory^33,52^.

Among the development-associated genes, we observe a diverse collection of relations between a gene’s expression and its chromatin activity, and importantly, the cross-modal relationship is clarified in Tilted-CCA’s common embedding. As expected, for many genes that are activated during development, their chromatin-level activity precedes their expression activation. Examples genes for this along the Glutamatergic 4 (Glu4) lineage in human embryonic brain, such as NTRK2, CNTN1, GRM1, and SEMASE, are shown in Figure 6a-c. This “priming” effect can be visualized as a time series across pseudotime, where chromatin activity increases gradually, preceding and foretelling the steep increase in gene expression much later (Fig. 6a). The phase portrait for the common components of RNA expression versus chromatin activity makes this relation more apparent with a chromatin-activity to RNA curve that “runs before rising” (Fig. 6b). Next, consider genes whose expression decreases over development, such as ASCL1, EOMES, PAX6 and ZNF521. We find that for these genes, chromatin activity drops much more gradually than RNA expression (Fig. 6d-f). The *cis*-peaks remain open after transcription of the gene has terminated, hinting at short-term cellular memory of the previous state. Lastly, and perhaps most curiously, we see evidence of genes that do not follow these aforementioned patterns but instead have a cyclical trend (Fig. 6g-i). Phase portraits for the other selected genes in Figure 5k are shown in Extended Data Figure 6c. In almost all cases, the relationship between RNA expression and chromatin activity is clarified in the Tilted-CCA common embedding, as shown by comparing to the insets of Figure 6b,e,h and the bottom rows of Figure 6c,f,i.

**Figure 6.**
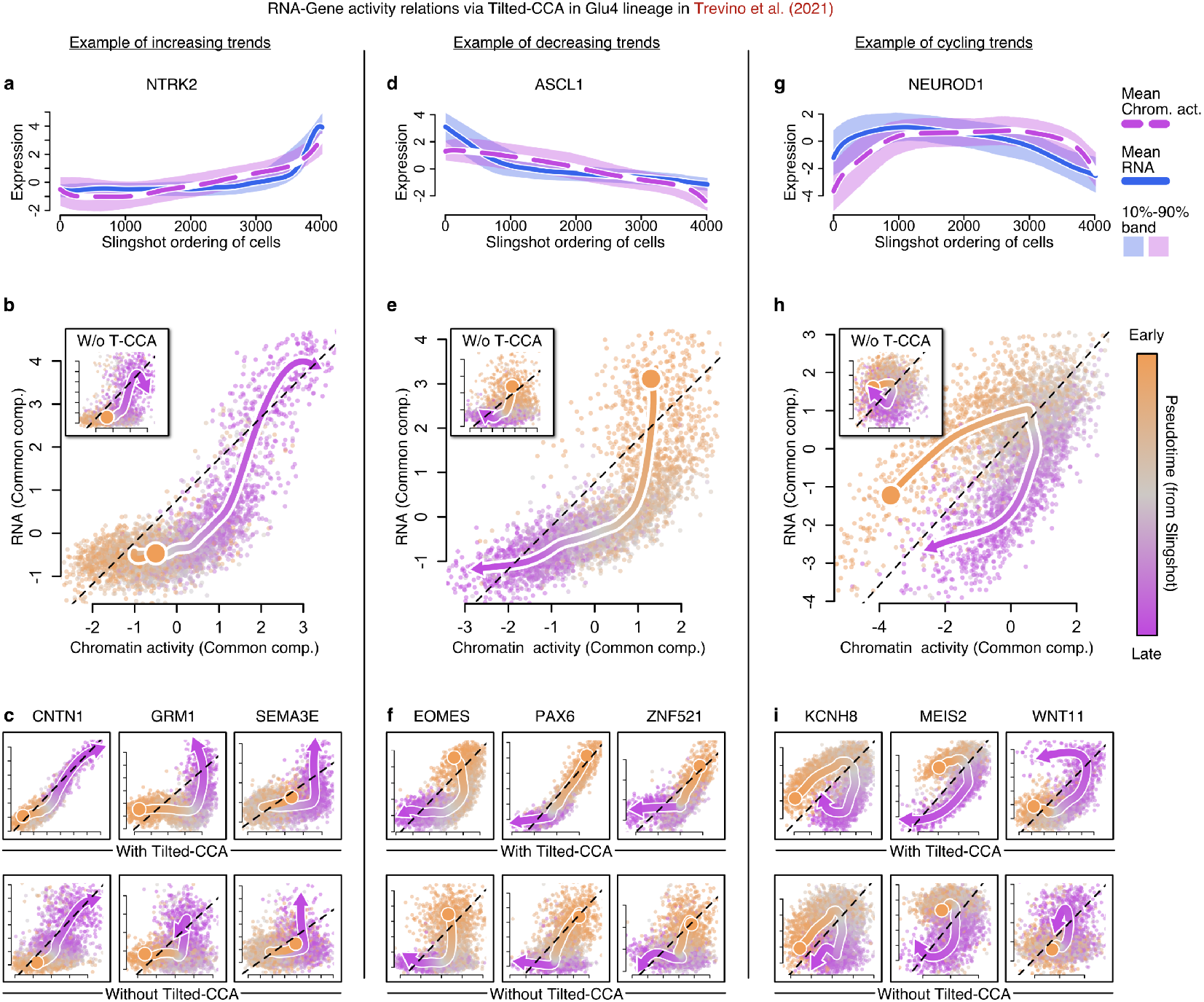
The relation between a gene’s expression and *cis*-chromatin activity is clarified by Tilted-CCA. **a,d,g**, Time-series showing the mean gene expression and cis-chromatin activity across pseudotime after applying Tilted-CCA for NTRK2, ASCL1 and NEUROD1 respectively, where each of the three genes illustrate different relations between the two modalities and all three genes are important for cortical neurogenesis. **b,e,h**, Phase portrait showing the mean gene expression plotted against *cis*-chromatin activity after for NTRK2, ASCL1 and NEUROD1 respectively applying Tilted-CCA, where cells are colored by the pseudotime. Each plot has a corresponding inset that shows the phase portrait if Tilted-CCA were not performed, illustrating noisier relationships between the two modalities. **c,f,i**, Additional phase portraits for other important genes among the three category of relations between gene expression and *cis*-chromatin activity, where the phase portraits are shown with (top) and without (bottom) applying Tilted-CCA.

## Discussion

Tilted-CCA extends CCA to decompose multimodal data into a sum of axes of variations that are shared between modalities and patterns that are distinct to each modality. We demonstrate the utility of Tilted-CCA on single-cell data, where we analyze multiomic datasets measuring the RNA and protein modalities or the RNA and ATAC modalities. For multimodal RNA and protein data, we show that Tilted-CCA aids in the design of targeted antibody panels that are complementary to the RNA modality. For multimodal RNA and ATAC data, we show how Tilted-CCA’s common embedding, which captures variation that is supported by both the RNA and ATAC modalities, can be used to estimate quantities related to cellular differentiation.

Tilted-CCA complements existing dimension-reduction methods for multiomic datasets because Tilted-CCA captures the intersection of information between modalities, while existing methods aim to capture the union of information. While these “union”-type methods, such as Consensus PCA, WNN, scAI, and MOFA+, are useful for aggregating information across modalities to improve cell-type discovery, these methods are not suitable for understanding how the two modalities are related to each another. Decomposition of multimodal data into common and distinct components is a fundamentally different question requiring new statistical methodology. Tilted-CCA answers this question, by first defining what is common information via the shared geometry of the two high-dimensional matrices, and then adapting the linear framework of CCA to best encapsulate this information. Importantly, this linear framework enables biologists to address either cell-centric or variable-centric questions in downstream analyses.

**Online content**

## Methods

### Mathematical foundation of Tilted-CCA

Let 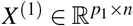 and 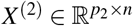 be the two matrices that form the multiomic dataset, where the same *n* cells are measured across *p*_1_ features in Modality 1 and *p*_2_ features in Modality 2. We assume that both *X*^(1)^ and *X*^(2)^ are preprocessed beforehand to have centered features. We also assume that both *X*^(1)^ and *X*^(2)^ are low-rank, i.e., rank(*X*^(1)^) = *r*_1_ and rank(*X*^(2)^) = *r*_2_ where *r*_1_ ≤ min{*n, p*_1_} and *r*_2_ ≤ min{*n, p*_2_}. See the remark near the end of the section about this low-rank assumption. Let *r* = min{*r*_1_, *r*_2_}.

Prior to describing the procedure, let us recapitulate the goal of Tilted-CCA from a mathematical perspective, which is to achieve the following decomposition:

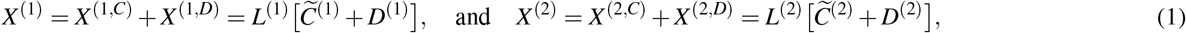

where the matrices 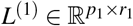 and 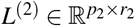 are modality-specific loading matrices

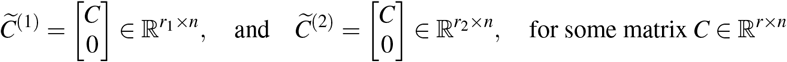

denote the common embeddings (where, in the situation of *r*_1_ = *r*_2_, then 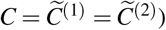, and 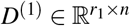 and 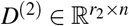 denote the modality-specific distinct embeddings. There are three constraints that we impose onto the decomposition in (1):

1. **Derivation from CCA**: The matrices 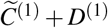 and 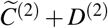 are the solution to CCA. Specifically, letting 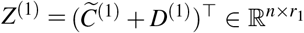 and 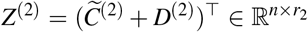, we require that the first *r* columns of *Z*^(1)^ and *Z*^(2)^ are the matrices of the leading r canonical score vectors. This ensures that the matrices *Z*^(1)^ and *Z*^(2)^ are suitable for decomposition where the first few columns are reflective of the the common embeddings (i.e., pairs of canonical score vectors that have high canonical correlation, capturing the “intersection of information”) and the last few columns are reflective of each modality’s distinct embedding (i.e., pairs of canonical score vectors that have low canonical correlation). Notably, this means that the matrices *Z*^(1)^ and *Z*^(2)^ are not designed for maximizing the predictive-power of the embedding, which is the goal of existing methods that capture the “union of information” instead (see Supplementary Information).
2. **Orthogonal distinct embeddings**: As mentioned in the in main text, we require

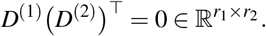 In combination with the first requirement, this mathematically formalizes what it means for the embeddings to be “distinct.”
3. **Approximation of common geometry**: Qualitatively, the common embedding *C* should capture geometry that is shared between *X*^(1)^ and *X*^(2) ”^ where, loosely-speaking, pairs of cells that are close in either modality should be close in C. This geometry is formalized by the target common manifold, described below, and can be thought as the population quantity to which Tilted-CCA is giving a linear approximation. From this perspective, the estimation of this (nonparameteric) population manifold through a linear decomposition of CCA’s canonical score vectors can be thought of as a form of linear regularization.

We now describe the procedure, with consists of five steps: (1) constructing nearest-neighbor graphs from each modality, (2) defining the target common manifold that encodes the common geometry between the two modalities, (3) performing CCA, (4) optimizing the appropriate tilt in the decomposition of CCA’s canonical score matrices such that the common space approximates the target common manifold, and (5) deriving the final decomposition in the form of (1).

#### Step 1: Construct nearest-neighbor graphs

The first step of Tilted-CCA is to construct two nearest-neighbor graphs, one for each *X*^(1)^ and *X*^(2)^. The procedure below has empirically worked well for our data examples, but mathematically, any reasonable nearest-neighbor graph construction would suffice. Let *k* denote the number of nearest neighbors, where *k* ≪ *n*. Let the SVD of *X*^(1)^ and *X*^(2)^ be denoted as

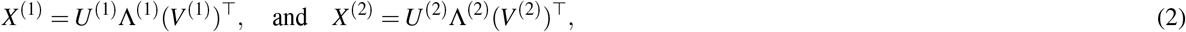

where, for both *∓* ∈ {1,2}, 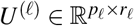 and 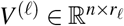 are orthonormal matrices, and 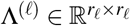 are diagonal matrices. Consider the low-dimensional embeddings 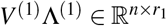 and 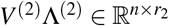. We normalize each cell’s embedding by constructing matrices 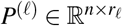 where

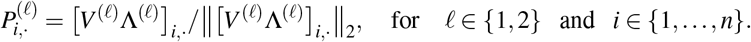

Then, we construct two nearest-neighbor graphs *G*^(1)^, *G*^(2)^ ∈ {0,1}^*n* × *n*^, where these graphs are represented as symmetric binary adjacency matrices among all *n* cells. Specifically, for *ℓ* ∈ {1, 2} and *i, j* ∈ {1,…, *n*} where *i* ≠ *j*

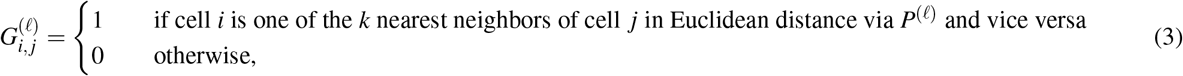

and 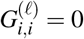 for all *i* ∈ {1,…, *n*}. Notice that computing Euclidean distances on *P*^(ℓ)^ is effectively computing cosine distances on the low-dimensional embedding *V*^(ℓ)^ *D*^(ℓ)^, and our requirement to have an edge if and only if cell *i* and *j* are both one of the *k* nearest neighbors of each other is often attributed to the shared nearest-neighbor (SNN) graph or mutual nearest-neighbor graph. Throughout rest of our explanation, we assume that *r*_1_ = *r*_2_ = *r* for simplicity of explanation, but the general procedure we’ve implemented allows for *r*_1_ ≠ *r*_2_, where *r* = min{*r*_1_, *r*_2_}.

#### Step 2: Construct target common manifold

The second step of Tilted-CCA is to determine the target common manifold based on the two nearest-neighbor graphs *G*^(1)^ and *G*^(2)^. This target common manifold *G* ∈ {0, 1}^*n*×*n*^ is itself also represented as a symmetric binary adjacency matrix. We have two different procedures for constructing G, based on whether or not a clustering is given for each of *X*^(1)^ and *X*^(2)^. For example, in developmental datasets, there is often a smooth continuum of cells in *X*^(1)^ and *X*^(2)^, meaning there is no well-defined clusters. On the other hand, in cell atlases where there different well-defined “hard” clusters of cells (possibly suggested by UMAPs), let

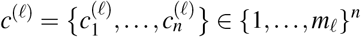

assign each of the *n* cells to *m_ℓ_* different clusters for *X*^(ℓ)^, for *ℓ* ∈{1, 2}. In our analyses, we use `Seurat::FindClusters` separately on *X*^(1)^ and *X*^(2)^ in this scenario. Importantly, typically *c*^(1)^ ≠ *c*^(2)^, since *X*^(1)^ and *X*^(2)^ have different clustering structure that will be relevant for constructing *G*. For either of these two cases, we have a different procedure to construct the target common manifold *G*. We have two separate procedures for each scenario since the former setting constructs *G* solely based on the local neighbor of each cell in *G*^(1)^ and *G*^(2)^, and the latter setting constructs *G* respecting the given clustering. The mathematical details are provided in the Supplementary Information, and we summarize the procedures below.

- **Scenario with no global clustering structure**: When no global structure is provided, the procedure to construct *G* is relatively straightforward. Intuitively, we include an edge between two cells in *G* if there is an edge present between the two cells in either *G*^(1)^ or *G*^(2)^. However, we perform a minor adjustment to ensure that both modalities contribute similar amounts of information towards *G*. For cell *i*, if Modality 1 and 2 have *a* and *b* unique edges involving cell *i* respectively where *a* > *b*, this involves purposefully downsampling the number of unique edges from Modality 1.
- **Scenario with global clustering structure**: When global structure is provided, the procedure to construct *G* is more involved, since we need to solve quadratic optimization program for each cell *i*. We require the user to specify *k* a priori, the desired number of edges per cell in *G*. (This is typically chosen to be the same *k* used to construct *G*^(1)^ and *G*^(2)^.) Broadly speaking, for cell *i*, the optimization program dictates how many edges from *G*^(1)^ or *G*^(2)^ are included in *G* so that both modalities contribute the same amount of information. This is mathematically quantified by first defining two multinomial distributions, one from each *G*^(1)^ and *G*^(2)^, that captures the proportion of edges connecting cell *i* to either set of hard clusters. Then, we use quadratic optimization to determine how to downsample from the total set of edges that yields a set of *k* edges that corresponds multinomial distributions closest to the two aforementioned multinomial distributions in *ℓ*_2_ distance.

Importantly, both constructions of *G* are randomized procedures, but we have found empirically that this source of randomness has little impact on the resulting Tilted-CCA embeddings.

#### Step 3: Perform CCA

The third step of Tilted-CCA is to perform CCA between *X*^(1)^ and *X*^(2)^. We review CCA here to lay out the notation. Let Σ^(1)^ = *X*^(1)^ (*X*^(1))⊤^ /*n*, Σ^(2)^ = *X*^(2)^ (*X*^(2)^)^⊤^ /*n* and Σ^(12)^ = *X*^(1)^ (*X*^(2)^)^τ^ /*n*. Recall that a rank-*r* CCA solves the optimization problem:

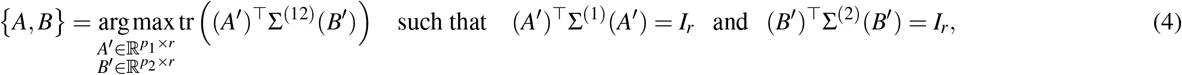

where *I_r_* is the *r*-dimensional identity matrix. There is an explicit closed-form solution to the optimization problem (4). Specifically, consider the SVD of the matrix

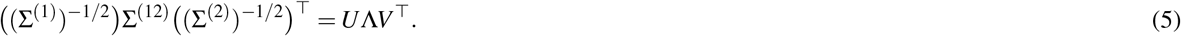

(It should be noted that if the value in the diagonal matrix *D* are in descending order, these diagonal values are the canonical values, i.e., values between 0 and 1 that denote the canonical correlation between the two modalities, where a value closer to 1 means there is more correlation.) (Σ^(1)^)^−1/2^ is the square-root matrix of Σ^(1)^. Here, because we are dealing with matrices *X*^(1)^ and *X*^(2)^ where potentially min{*p*_1_, *p*_2_} ≥ *r*, we define the square-root matrix not as a square matrix. Instead, recalling the SVDs in (2), we define for *ℓ* ∈{1, 2},

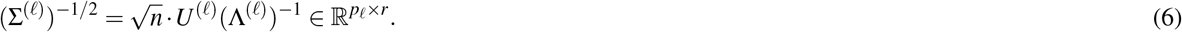

With this definition of the square-root matrix, observe that (Σ^(*ℓ*)^)^−/2^ Σ^(*ℓ*)^((Σ^(*ℓ*)^)^−1/2^)^⊤^ = *I_r_*. Notice that since the rank of Σ^(12)^ is at most *r*, we see that 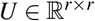 and 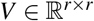 in (5). The solution to (4) is then

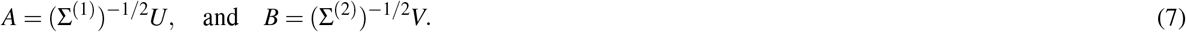

Equipped with the solution to (4) given in (7), the SVDs given in (2) and the square-root matrices in (6), we can compute the canonical score matrices for *ℓ* ∈{1, 2},

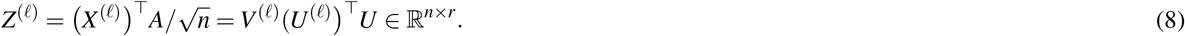

Note that by the identity constraints of CCA in (4), *Z*^*(ℓ*)^ is an orthonormal matrix, i.e., 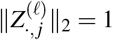 for all *j* ∈ {1,…, *r*}. (We can verify this holds since 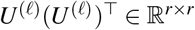 is the identity matrix – it projects vectors in a vector space of dimension *r* onto the full-rank basis of a vector space of dimension *r*.)

#### Step 4: Optimize the common and distinct scores

The fourth step is to compute the common and distinct scores, using the target common manifold and the canonical scores *Z*^(1)^ and *Z*^(2)^. Let 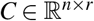 denote the common score matrix, and 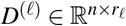 denote the distinct score matrix for Modality *ℓ* ∈ {1, 2}. Specifically, we require that for *ℓ* ∈ {1, 2},

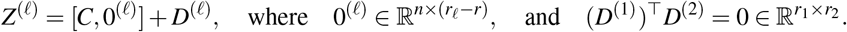

Next, to construct *C, D*^(1)^ and *D*^(2)^ to satisfy these requirements, we describe how a proposed tilt enables us to construct these three matrices. The construction is done column-wise, for latent dimension *j* ∈ {1,…, *r*}. Consider a tilt *τ_j_* ∈ [0,1], where if *τ_j_* = 0 then 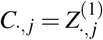 and if *τ_j_* = 1 then 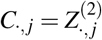. Specifically, there is a unique construction of 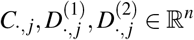 based on 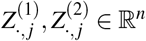 and *τ_j_* ∈ [0,1], such that

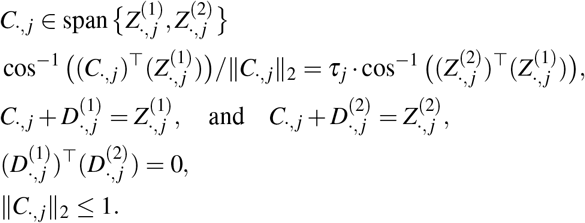

The first relation ensures that we are only considering common vectors in the same hyperplane as the two canonical score vectors – this ensures that the resulting column vectors *C* are orthogonal after this construction is complete. The second relation is why we call *τ_j_* the “tilt” – we are ensured that *C*_.,*j*_ is a vector that has an angle of *τ_j_*-percent between 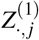 and 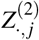. The third and fourth relations ensure a valid decomposition of 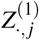 and 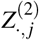. The fifth equality relation the two distinct vectors 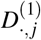 and 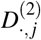 are orthogonal, which is why we call these the “distinct” vectors. The last relation ensures a unique decomposition. The vectors *C*_.,*j*_, 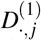 and 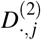 can be constructed to satisfy these constraints using straightforward geometry and linear algebra (See Figure1h for the intuition on the details of the calculation).

Lastly, we describe how we measure the quality of a proposed tilt *τ*_*j*_ based on its similarity to the target common manifold *G* constructed in the second step. Consider the common manifold *G* described in Step 2 and the nearest-neighbor graph constructed from *C* with tilt *τ*_*j*_, denoted as *G*^(*C*;*τ*_*j*_)^. (Specifically, we apply the same construction in (3), but based on *C* instead of *P*^(ℓ)^.) Equipped with *G* and *G*^(*C*;*τ*_*j*_)^, our similarity is defined by the Grassmannian distance between two sets of eigenvectors, each derived from the normalized random-walk Laplacian graph basis^53^ for either of these graphs. Specifically, considering the common manifold *G*, let 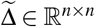 denote the degree matrix which is a diagonal matrix where

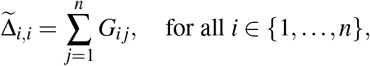

and define the normalized Laplacian as 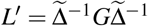, and then define the normalized random-walk Laplacian as

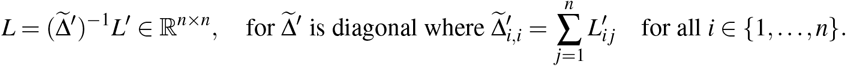

Using this construction, let *W* and *W*^(*C*;*τ_j_*)^ be the leading-*K_L_* eigenvectors of *L* and *L*^(*C*;*τ_j_*)^ normalized random-walk Laplacian basis matrices constructed from the graphs *G* and *G*^(*C*;*τ*_j_^) respectively, for some tuning parameter *K_L_*. Since both *W* and *W*^(*C*;*τ_j_*)^ are points along the Grassmannian manifold, we use the Grassmannian distance^54,55^ to measure the distance between *W* and *W*^(*C*;*τ_j_*)^. Specifically, consider the SVD of

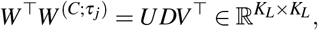

where *D* = diag(σ_1_,…, *σ_K_L__*). The Grassmannian distance is then defined as

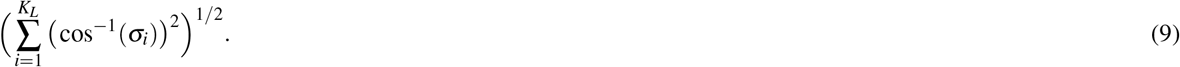

A smaller Grassmannian distance implies a higher similarity between *G* and *G*^(*C*;*τ*_j_)^.

We now have all the necessary ingredients – we have described how to construct *C* (and hence, *D*^(1)^ and *D*^(2)^) based on a proposed tilt *τ_j_*, one tilt for each of the *r* latent dimensions, and we also know how to measure the quality of the proposed tilted using the aforementioned Grassmannian distance. All that remains is an optimization procedure. Towards this end, we use a zero-order cyclical (either using Nelder–Mead or a grid-search) optimization over *τ_j_* ∈ [0,1] each latent dimension *j* ∈ {1,…, *r*} to minimize the distance between the target manifold *G* and the common embedding *G*^(*C*;*τ_j_*)^ and we cycle through each latent dimension iteratively (i.e., many epochs) until we reach convergence or an maximal epoch limit.

#### Step 5: Compute the final decomposition

The last step is to determine the decomposition of *X*^(1)^ and *X*^(2)^ based on the common and distinct scores. Specifically, for Modality *ℓ* ∈{1, 2}, let

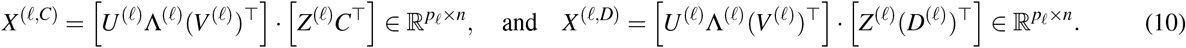

This is a sensible construction, as 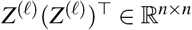 is a projection matrix since 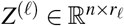 is an orthonormal matrix and *C* + *D*^(*ℓ*)^ = *Z*^*ℓ*^, while *U*^*ℓ*^ ∧^(*ℓ*)^(*V*^(*ℓ*)^)^⊤^ is each modality’s low-rank expression matrix. Hence, by this construction, we are ensured that

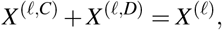

which justifies why this is a decomposition, and that *X*^(1, *C*)^ and *X*^(2, *C*)^ share the same row-space, which justifies why this is called the common component of the decomposition.

Observe that from the form of (1), the loading matrices can be analogously calculated as

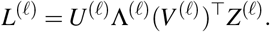

##### Remark about computation

The above procedure can be computationally-exhaustive when the number of cells *n* is more than 10,000. This stems from the fact when optimizing for *τ_j_* (for each latent dimension *j* ∈ {1,…, *r*} over a couple epochs), many graphs *G*^(*C*;*τ_j_*)^ (the nearest-neighbor graph of the common embedding for a posited *τ_j_* for latent dimension *j*) need to be computed. Constructing these nearest-neighbor graphs can be quite expensive for large *n*’s. Our strategy to reduce the computational cost of Tilted-CCA is to compute nearest-neighbor graphs on *n_m_* meta-cells (where *n_m_*≪ *n*) instead of on the original *n* cells.

Specifically, suppose we have a hard-clustering of the *n* cells into *n_m_* meta-cells, and let *Ā* ∈ [0, 1]*^n_m_ × n^* be the averaging operator where for any matrix 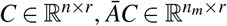 is the matrix where the *i*th row is constructed by row-wise averaging of *C* corresponding to the cells (i.e., columns) in the ith meta-cell. This can be constructed in a variety of ways, of which we describe the particular strategy we use later. Equipped with this meta-cell, we can tweak the Tilted-CCA procedure to accommodate the meta-cells in the following way:

- The nearest-neighbor matrices *G*^(1)^ and *G*^(2)^ are constructed based on the columns of 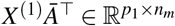 and 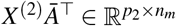, the average expression matrices of each meta-cell for each modality. This means *G*^(1)^ and *G*^(2)^ are both matrices of dimensionality *n_m_* × *n_m_*.
- Since *G*^(1)^, *G*^(2)^ ∈{0, 1}*^n_m_^ ×^n_m_^*, when we construct the target common manifold *G*, it is also of dimensionality *n_m_* × *n_m_*.
- The CCA is performed on the original *n* cells, and when optimizing for the tilt *τ_j_* for latent dimension *j* ∈ {1,…, *r*}, the zero-order grid search yields a common embedding 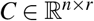 for a posited *τ* ∈ [0, 1]. Then, we construct its associated nearest-neighbor graph based on the rows of *ĀC*, yielding *G*^(*C*;*τ_j_*)^ ∈ {0, 1}*^n_m_ × n_m_^*. Then, the Grassmannian distance (9) is computed between *G* and *G*^(*C*;*τ_j_*)^.

We propose the following way to obtain the meta-cells (although many other reasonable procedures can be used). Supposing we desire *n_m_* meta-cells, we partition the *n* cells in each modality into 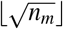 clusters (for example, using K-means on the PCA embedding of X^*ℓ*^ for Modality *ℓ* ∈ {1, 2}). This yields two clusterings, 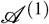 and 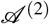. Then, we intersect the 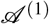 and 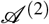 to yield 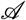, a clustering with at most *n_m_* clusters (meaning two cells are in the same cluster in 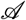 if and only if both cells are in the same cluster in both 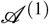 and 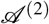). If 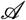 has less than *n_m_* clusterings, we further partition each cluster in 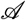 using K-means based on the size of the clusters in 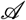 to yield a total of *n_m_* clusters.

##### Remark about tuning the parameters

To summarize, the tuning parameters that a practitioner would need to choose prior to applying Tilted-CCA are as follows:

- Parameters relevant to constructing each modality’s nearest-neighbor graph (3). This primarily involves choosing the number of nearest neighbors *k*, as well as the dimensionality *K_L_* of the graph Laplacian basis for the graphs *G*^(*C*;*τ_j_*^) and *G*. These parameters will be used to construct the target common manifold.
- The number of latent dimensions for each modality. Recall that in full generality, Tilted-CCA allows the usage of the leading *r*_1_ and *r*_2_ canonical vectors from Modality 1 and 2 respectively, where *r*_1_ might not necessarily be equal to *r*_2_. In practice, we defer to the typical preprocessing pipelines to appropriately choose the appropriate number of latent dimensions for each modality. For example, this could be done using scree plots, jackstraw methods, visualizations of the derived low-dimensional embedding, or other statistically-driven methods.
- Potential global clustering structure when designing the target common manifold, if the practitioner observes visibly-separable separate among cell-types in one of the two modalities.
- Additional parameters relevant to the optimization procedure or meta-cell construction. These include the termination threshold to stop the cyclical optimization procedure for the tilt of the common vectors, as well as number of meta-cells (if the practitioner desires to use meta-cells to ease the computational burden).

In practice, the least-obvious parameter to choose is the number of nearest neighbors *k* and the dimensionality of the graph Laplacian bases *K_L_* for the nearest-neighbor graph constructions. We suggest the following diagnostic to assess if an appropriate value of *k* is chosen: first, for a particular value of *k*, compute each modality’s nearest-neighbor graph, *G*^(1)^ and *G*^(2)^. Then, this enables one to derive the target common manifold *G* using the same tuning parameter *k*. Then, visualize *G* (for example, by plotting a UMAP or a rank-2 diffusion map of the leading-*K_L_* graph Laplacian eigenbases of *G*). By comparing this visualization to the visualizations of Modality 1 and 2 (for example, UMAPs of *X*^(1)^ and *X*^(2)^), the practitioner can assess if the target common manifold is appropriately capturing the “intersection” of information – does it visually seem apparent that cells that separable in both modalities are separable in G? Through this visual diagnostic, the practitioner would have more assurance that Tilted-CCA’s optimization procedure yields a linear decomposition that is approximating an appropriate common manifold. After the entire Tilted-CCA procedure is done, the enrichment plots analogous to those in Extended Data Figure 2a,b can be used to diagnose if the common embedding captures biological expectations. These diagnostic checks are vital before proceeding to any of the downstream analyses performed in this paper, as they rely on an appropriately-estimated common and distinct embeddings.

### Quantifying enrichment of a cell type

We describe the details of how to compute the enrichment of a cell type in the distinct embedding (i.e., based on *D*^(1)^ or *D*^(2)^) in order to assess which modality better separates which cell types, illustrated in Figure 3b and Extended Data Figure 2a,b. We first describe the intuition behind our method, and then describe the mathematical details. We first construct a SNN graph based on either modality’s distinct embedding. Then, given an external labeling of which of the *n* cells belong to which cell type, for a particular cell-type, we compute the average proportion of each cell having nearest neighbors of the same cell type, adjusted for the baseline proportion of said cell type. (Recall, Tilted-CCA is an unsupervised method, so these cell-type labelings are not used to compute the distinct embeddings themselves.) Mathematically, for Modality *ℓ* ∈{1, 2} we first calculate the matrix,

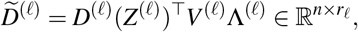

which represents the distinct embedding with columns reweighted in accordance to the prevalence of the corresponding features in the modality *X*^*ℓ*^. Note that this construction is similar to that in (10) – here, 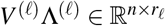 represents the modality’s low-dimensional embedding.

Using 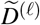, we then compute the SNN graph *G*^(*ℓ*;*D*)^ ∈{0, 1}^*n*^ ×^*n*^ using the same procedure as described in the “Step 1: Construct nearest-neighbor graphs” subsection above. Then, for a particular cell type, consider the set of cells of that cell-type, denoted as the index set 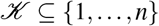 where 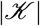 denotes the cardinality of that set. Then, for cell 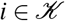, let 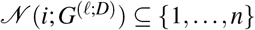 denote the neighbors of cell *i* in graph *G*^(*ℓ*;*D*)^. The enrichment of this cell-type is then computed as

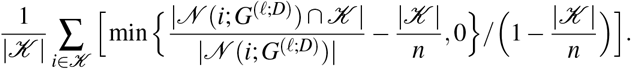

Observe this is the average proportion among all 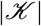 cells of the cell type, where the proportion is adjusted for the baseline proportion by subtracting 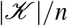 from the numerator and denominator. This is justified since a cell type prevalent in the dataset will naturally be neighbors of many cells.

Observe that this procedure is not limited to the distinct modalities of Tilted-CCA. The above procedure relies on having a nearest-neighbor graph and cell-type labelings among the *n* cells. Hence, it could also be applied to the common embedding, or the Consensus PCA embedding illustrated in Figure 4e.

### Constructing alignment-differentiability scatterplots

We describe the details of how to construct the alignment-differentiability scatterplots in order to assess which features have expression patterns reflected in other modality, illustrated in Figure 3c-e, Extended Data Figure 2c,e, and Supplementary Figure 2. These plots rely on computing the alignment score and differentiablity score for each feature. We describe each in detail, where we assume that procedure is applied to each gene for simplicity, where the RNA modality is *ℓ* = 1.

- **Alignment**: For each gene *j* ∈ {1,…, *p*_1_}, compute a linear regression that regresses 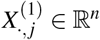 onto 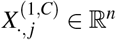, with an intercept term. Then, return the *R*^2^ of this regression.
- **Differentiability**: Let 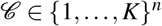 denote the externally-provided cell-type labels for the *K* cell types among the *n* cells. For each gene *j* ∈ {1,…, *p*_1_} and for each pair of cell-types *k*_1_, *k*_2_ ∈ {1,…, *K*}, compute the two-sample test of difference in means between all cells of cell-type *k*_1_ and cells of cell-type *k*_2_ based on *X*^(1)^ for gene *j*, and let *ρ*^(*j*;*k*_1_,*k*_2_^) denote the p-value of this hypothesis test. Since we are applying this procedure to the RNA modality, we use MAST^56^ test since this accounts for the library size of the cells. (For differential expression tests among antibodies, we default to the Wilcoxon rank-sum test.) Then, we summarize all such *ρ*^(*j;k*_1_,*k*_2_)^ values for a particular cell-type *k*_1_ in the following way:

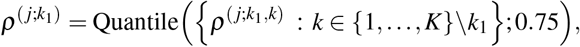

which denotes the 75th quantile among all *ρ*^(*j*;*k*_1_,*k*_2_)^ values involving cell-type *k_1_*. Finally, for gene *j*, return

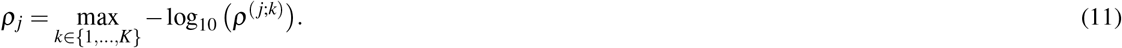

The alignment score is designed in this way as a gene is highly alignable if its common component contains similar expression patterns as the observed gene expression itself. The differentiabilty score is designed this way since a gene is highly differentiable if it separates a few particular cell type from most other cell types.

Observe that this procedure is not limited to the RNA modality. It could applied to antibodies in the protein modality or the chromatin regions in the ATAC modality.

### Designing antibody panel panel using Tilted-CCA

We describe the details of how to generate targeted antibody panels that contain differential-expressed signals among surface antibody markers designed to compliment the RNA modality for RNA+AB platforms such as CITE-seq and Abseq, illustrated in Figure 4 and Extended Data Figure 4. Intuitively, the procedure first computes the Tilted-CCA and then computes how differentially-expressed the distinct components of an antibody expression is. Then, the method performs greedy selection – it finds the first antibody that with the highest differentially-expressed distinct component whose total expression is uncorrelated with the common subspace estimated by Tilted-CCA, adds that antibody to a growing set of selected antibodies, and then continues iteratively by finding another antibody with the highest differentially-expressed distinct component among the remaining antibodies whose total expression is uncorrelated with both the common subspace and the already-selected antibodies.

Mathematically, assuming the protein modality is *ℓ* = 2, let *r*^(thres)^ ∈ [0,1] be a user-chosen correlation threshold (a suitable default being *r*^(thres)^ = 0.85), and *P* < p_2_ be the user-chosen antibody panel size (a suitable default being *P* = 10). The remaining inputs to our procedure are: 1) the antibody expression matrix, 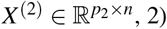, 2) the differentiabilty of each antibody’s distinct component, measured on the –log_10_ scale, *ρ*_1_,…, *ρ*__*p*2__ ≥ 0, akin to our differentiability score in (11) but computed based on *X*^(2, *D*)^, and 3) the common embedding given by Tilted-CCA, 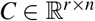, which we initialize the matrix 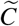 to be. Then, the procedure follows the iterative procedure, after initializing the set of candidate antibodies to be 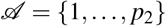 and the selected antibody panel 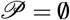.

1. Select all antibodies in 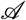 whose expression vector in *X*^(2)^ has an *R*^2^ less than *r*^(thres)^ when regressed onto 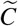 (involving an intercept term). This regression is performed on each antibody separately. Let this set be the new 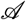. Terminate if 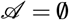.
2. Among all the antibodies in 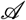, add the antibody with largest *ρ_j_* (i.e., differentiability among its distinct component) to the panel 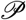, and then include its expression vector 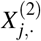 to 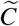.
3. If the size of 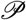 is less than 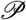, continue. If not, return 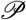.

By selecting antibodies by regressing *X*^(2)^ onto 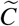 (instead of regressing *X*^(2;*D*)^ onto 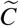, we naturally account for how substantial an antibody’s distinct component is relative to its common component.

### Selecting development-associated genes

We describe the details of how to select development-associated genes, illustrated in Figure 5e,g and and Extended Data Figure 6b. Intuitively, the procedure first regress each gene’s RNA expression as well as its chromatin activity onto the graph Laplacian basis of Tilted-CCA’s common embedding (formed jointly between the RNA and chromatin activity modalities). Then, the method performs greedy selection – it first finds the gene with the highest *R*^2^ in both the RNA and chromatin activity modalities, adds that gene to a growing set of development-associated genes, and then continues iteratively by finding another gene with the highest *R*^2^ whose RNA and chromatin activity modalities are sufficiently uncorrelated with previously-selected genes. This selection procedure bears strong resemblance to the method to select antibody panels.

Mathematically, let *ρ*^(thres)^ ∈ [0,1] be a user-chosen correlation threshold and *P* be the user-chosen number of development-associated genes. The remaining inputs to our procedure are: 1) *X*^(1)^, 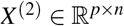 denote the RNA and chromatin activity modalities respectively, 2) the graph Laplacian for Tilted-CCA’s common embedding 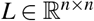, and 3) two sets of *R*^2^ vectors, 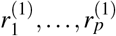 and 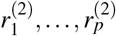 either denoting the *R*^2^ of regressing each gene in *X*^(1)^ or *X*^(2)^ onto *L* respectively (including an intercept term). Then, select the first gene

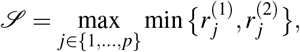

as in other words, the maxi-min gene measured by *R*^2^. Then procedure follows the iterative procedure, after initializing the set of candidate genes to be 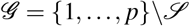.

- Select all genes in 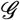 satisfying: 1) its expression vector in *X*^(1)^ is correlated less than *ρ*^(thres)^ to the expression vector in *X*^(1)^ of any already-selected gene in 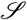, and 2) its expression vector in *X*^(2)^ is correlated less than *ρ*^(thres)^ to the expression vector in *X*^(2)^ of any already-selected gene in 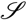. Let this set be the new 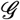. Terminate if 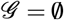.
- Among all the genes in 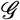, add the gene with largest minimum (between 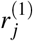 and 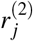) to the set 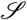.
- If the size of 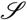 is less than *P*, continue. If not, return 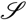.

To clarify, this procedure as described does not necessarily involve the Tilted-CCA as any reasonably-chosen graph that encapsulates the trajectory geometry can be used, such as the SNN graph from only the RNA modality). Nonetheless, 1) we’ve found the SNN graph of the common embedding to be more stable than the SNN graph based on Consensus PCA, WNN, or either the RNA or chromatin activity modality only as the SNN graph of the common embedding is corroborated by both modalities, and 2) we believe there are benefits of such a procedure compared to existing gene-selection methods since the biologist does not need to commit to a particular pseudotime ordering of the cells prior to selecting the development-associated genes. Also, the procedure does not need to be necessarily done jointly between the RNA and chromatin activity modalities in general, but we have found the quality of the genes to be noticeably more appropriate compared to when we selected genes based on only the RNA expressions.

### Computing the synchrony score between RNA and ATAC

We describe the details of how to compute the synchrony scores of a cell’s genes based on its peaks in order to infer if a cell is in steady-state, illustrated in Figure 5b,d,f and Extended Data Figure 6a. We first describe the intuition behind our method, and then describe the mathematical details. Recall that we posit that the common subspace between the RNA and ATAC is a suitable subspace that shows the developmental trajectory – this is because it is biologically expected that the developmental trajectory is reflected by the gene expression dynamics as well as the epigenetic dynamics from the opening/closing of chromosomes. Hence, our method’s goal is to assess if the distinct RNA and distinct ATAC expression patterns for a particular cell is both large in magnitude and orthogonal to its common expression pattern. If so, then the cell is likely not in steady-state, as either its distinct RNA or ATAC component contains information relevant to development that is not already encapsulated in the common subspace. The challenge of designing a method to accomplish this goal is that the ATAC and RNA modalities have differing number of features and possibly have expression values on different scales.

To accomplish this, our high-level idea is to first compute each a suitable linear transformation to rescale/rotate the low-rank ATAC modality to match the low-rank RNA modality, and then compute the correlation between this transformed ATAC expression vector and the common RNA expression vector for each cell. Two things are noteworthy:

- First, we are interested in common RNA expression vector as opposed to the common ATAC expression vector since the former has features that are more interpretable (i.e., genes).
- Second, by linearly transforming the low-rank ATAC modality to match the low-rank RNA modality, we are inherently accounting for both the distinct RNA and distinct ATAC components, relative to their common counterparts.

Mathematically, let 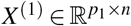 and 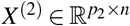 denote the low-rank RNA and ATAC modalities respectively (for *p*_2_ » *p*_1_ typically), where we seek to linearly-transform *X*^(2)^ to qualitatively match *X*^(1)^, that is, to find a low-rank transformation matrix 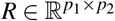 such that

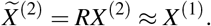

Formally, since both modalities have differing number of features yet are low-rank, this transformation is done through the right singular vectors of each modality. Denote the SVDs of both *X*^(1)^ and *X*^(2)^ as

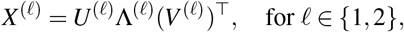

and assume for simplicity that both matrices are of rank *r* ≪ min{*n, p*_1_, *p*_2_}. Here, 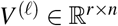. Then, recalling the solution to the Orthogonal Procrustes problem,^57,58^, we compute

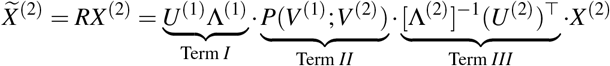

where Term *I* is a matrix of dimension *p*_1_ × *r* and Term *II* is a matrix of dimension *r* × *r* obtained as the solution of the Orthogonal Procrustes problem, and Term *III* is a matrix of dimension *r* × *p*_2_. Briefly, the Orthogonal Procrustes problem computes a matrix 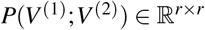 that optimally rotates the orthonormal matrix *V*^(2)^ to have the smallest difference from *V*^(1)^ measured in Forbenius norm. This matrix is computed by first computing the SVD of the following matrix,

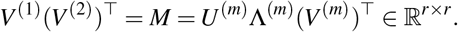

Using this notation, the solution to the Orthogonal Procrustes problem is

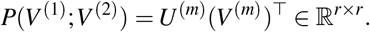

All together, from right to left, Term *III* whitens the ATAC modality to become “unitless” as [∧^(2)^]^−1^(*U*^(2)^)^⊤^*X*^(2)^ = *V*^(2)^, the right singular vectors of the ATAC modality. Term *II* then optimality rotates *V*^(2)^ to match *V*^(1)^, the right singular vectors of the RNA modality. Lastly, Term *I* transforms the resulting matrix to have one variable per gene and have values on the same scale as the RNA modality. Observe that 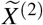 can be computed without fitting Tilted-CCA. Note that this procedure is not numerically equivalent to performing linear regressions since we want to predict the RNA modality from the ATAC modality while utilizing of the low-rank nature of both matrices.

Lastly, to finalize our predictability score between the RNA and ATAC modalities for each cell, we compute,

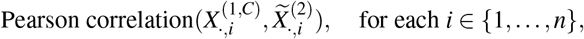

where 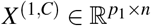 is the common RNA component resulting from Tilted-CCA, as computed in (10). Here, both vectors involved in the Pearson correlation are of length *p*_1_ (i.e,. the number of genes). A higher Pearson correlation qualitatively means that the RNA modality is more predictable from the ATAC modality, meaning the distinct components in either modality are either small in magnitude or have similar expression patterns as those in the common subspace.

We also note that the plots in Figure 5a were generated by plotting the absolute relative residual, i.e., 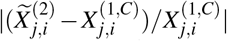, against Slingshot’s estimated pseudotime for cell i among all the cells along the Glutamatergic 4 lineage, for a particular gene *j*.

### Data processing of multiomic data

We describe the high-level procedure for analyzing the multiomic datasets. As assumed by Tilted-CCA, each modality is first preprocessed separately to be low-rank prior to being analyzed by Tilted-CCA. The preprocessing procedure should match the generally-accepted practices of the field, of which we describe the preprocessing we used for the RNA, protein, ATAC and chromatin activity modalities.

- RNA modality: Our preprocessing strategy of the scRNA-seq modality follows the procedure performed by the original authors of the dataset. This broadly dichotomizes into two categories. In the first category, the typical `Seurat` pipeline is used where the data matrix is first normalized via `Seurat::NormalizeData` via log-normalization, then highly-variable genes are first selected, followed by recentering and rescaling each of the highly-variable genes across all cells and projecting the data onto the leading principal components. In the second category, the data matrix is normalized via `Seurat::SCTransform`, which also selects the highly-variable genes. The genes are then recentered and rescaled across all the cells and projecting the data onto the leading principal components.
- Protein modality: We preprocess the protein modality analogous to the RNA modality via the first category (i.e., using log-normalization).
- ATAC modality: We preprocess the ATAC modality using the suggest `Signac` pipeline. Specifically, the highly-variable peaks are first selected via `Signac::FindTopFeatures` using `min.cutoff=‘q0’`. Then, among the selected peaks, data matrix is normalized via the TF-IDF transformation, and then projected onto the leading singular vectors via `Signac::RunSVD`. Two aspects are note-worthy: 1) As suggested by typical Signac pipelines, the first singular vector is omitted during the projection. 2) Due to the extreme sparsity typical of ATAC data matrices, the expression values are not rescaled or recentered. However, following the implementation of `Signac::RunSVD`, we often recenter/rescale the resulting singular vectors whenever appropriate in Tilted-CCA when working with the ATAC modality’s axes of variation.
- Chromatin activity modality: To compute the chromatin activity matrix, we apply `Signac::GeneActivity` where we sum the peak expressions ±500 bases from each gene’s TSS, which `biotypes=“protein_coding”`. Then, we normalized the resulting data matrix via `Seurat::NormalizeData` via log-normalization and set the “highly-variable genes” to be RNA modality’s highly-variable genes. These genes are then recentered and rescaled across all the cells and projecting the data onto the leading principal components, omitting the first principal component.

Here are additional relevant details for specific datasets.

- Human bone marrow CITE-seq (RNA+protein)^18^ : As suggested by the authors, the RNA modality was normalized via log-normalization and we used the leading 30 principal components for the RNA modality and the leading 18 principal components for the protein modality. We found *k* = 30 and *K_L_* = 20 to be suitable for analyzing this dataset using Tilted-CCA, and used global clustering structure estimated via `Seurat::FindClusters` with `resolution=0.25` for each modality, and used 5000 metacells when optimizing the tilts. The 25 antibody markers in this dataset are: CD11a, CD11c, CD1234, CD127-IL7Ra, CD14, CD16, CD161, CD19, CD197-CCR7, CD25, CD27, CD278-ICOS, CD28, CD3, CD34, CD38, CD4, CD45RA, CD45RO, CD56, CD57, CD69, Cd79b, CD8a, HLA.DR.
- Human bone marrow Abseq (RNA+protein)^20^ : There two datasets, both with 97 antibody markers. They differed in the RNA modality, one where only 461 genes were sequenced (a dataset with more cells) and another that sequenced the whole transcriptome (with less cells), and both datasets were already normalized when provided. Analogous to the authors’ analysis via MOFA+, we used the leading 30 principal components for the RNA modality and the leading 30 principal components for the protein modality. For the former dataset, we found *k* = 15 and *K_L_* = 20 to be suitable for the analyzing using Tilted-CCA where we used global clustering structure estimated via `Seurat::FindClusters` with `resolution=0.25` for each modality. For the latter dataset, we used *k* = 30 and *K_L_* = 20 and the global clustering structure estimated via `Seurat::FindClusters` with `resolution=0.1` for each modality. For both datasets, we used 5000 metacells when optimizing the tilts.
- Human PBMC CITE-seq (RNA+protein)^11^ : The data was normalized when provided. As suggested by the authors, we used the leading 40 principal components for the RNA modality and the leading 50 principal components for the protein modality. We found *k* = 30 and *K_L_* = 30 to be suitable for analyzing this dataset using Tilted-CCA, used global clustering structure estimated via `Seurat::FindClusters` with `resolution=0.25` for each modality, and used 5000 metacells when optimizing the tilts.
- Human PBMC 10× Multiome (RNA+ATAC): Following the suggested preprocessing pipeline in the Signac vignette to process this data, we first screened cells to have a total ATAC count between 500 and 7000 and a total RNA count between 1000 and 25000 and have a percentage of counts from mitochondrial genes to be less than 20%. Then, the leading 50 principal components for the RNA and ATAC modalities were used. We found *k* = 60 and *K_L_* = 50 to be suitable for analyzing this dataset using Tilted-CCA, used global clustering structure estimated via `Seurat::FindClusters` with `resolution=0.25` for each modality, and used 5000 metacells when optimizing the tilts.
- Human developing brain 10× Multiome (RNA+ATAC)^40^ : The leading 50 principal components for the RNA and ATAC modalities were used. In the analysis of RNA and ATAC, we found *k* = 15 and *K_L_* = 20 to be suitable for analyzing this dataset using Tilted-CCA, where no global clustering structure nor metacells were used. The same parameters were used when analyzing the RNA and chromatin activity.
- Mouse developing brain 10× Multiome (RNA+ATAC): The leading 50 principal components for the RNA and ATAC and chromatin activity modalities were used. In the analysis of RNA and ATAC, we found *k* = 30 and *K_L_* = 20 to be suitable for analyzing this dataset using Tilted-CCA. In the analysis of RNA and ATAC, we found *k* = 20 and *K*_L_ = 60 to be suitable. No global clustering structure nor metacells were used in either analysis.
- Mouse developing embryo 10× Multiome (RNA+ATAC)^49^ : The leading 50 principal components for the RNA and ATAC modalities were used. We found *k* = 30 and *K_L_* = 20 to be suitable for analyzing this dataset using Tilted-CCA. No global clustering structure was used and the optimization of the tilts were done using 5000 metacells.

## Data availability

All data used in this manuscript are publicly available. Specifically, the human bone marrow CITE-seq data^18^ is available in the R package `SeuratData` under “`bmcite`”. The human bone marrow Abseq data^20^ (both, either equipped with the whole transcriptome or of only 461 genes) is availabe at https://figshare.com/projects/Single-cell_proteo-genomic_reference_maps_of_the_human_hematopoietic_system/94469, under WTA_projected.rds or Healthy.rds respectively. The human PBMC CITE-seq data^11^ is available at https://atlas.fredhutch.org/nygc/multimodal-pbmc/. The human PBMC 10× Multiome data is available at https://support.10xgenomics.com/single-cell-multiome-atac-gex/datasets/1.0.0/pbmc_granulocyte_sorted_10k. The human brain development 10× Multiome data^40^ is available at https://www.ncbi.nlm.nih.gov/geo/query/acc.cgi?acc=GSE162170 and https://github.com/GreenleafLab/brainchromatin. The mouse brain development 10× Multiome data is available at https://www.10xgenomics.com/resources/datasets/fresh-embryonic-e-18. The mouse embryo development 10× Multiome data^49^ is available at https://github.com/rargelaguet/mouse_organogenesis_10x_multiome_publication. The majority of the computational tools used in this manuscript consisted of `Seurat` (v4.1.1), `Signac` (v1.7.0), and `slingshot` (v2.3.1). We derived the cell-cycle genes for humans via Seurat::cc.genes, while we used the lists of human housekeeping genes^59^ or mouse cell-cycling genes^60^ from particular published work.

## Code availability

The Tilted-CCA software package is available at https://github.com/linnykos/tiltedCCA. The scripts to reproduce the processing, analysis and figure-generation are available at https://github.com/linnykos/tiltedCCA_analysis.

## Acknowledgements

We thank Z. Ma, M. Li, CY. Wu, P. Hess, D. Matthew, K. Roeder and B. Devlin for helpful comments, suggestions, and feedback on the method development and analyses.

## Author contributions statement

K. Z. L. implemented the method and performed the computational analyses. K. Z. L. and N. R. Z. designed the method and suite of experimental results, and wrote the manuscript.

## Additional information

**Extended data.** See Extended Data Figures 1 through 6.

**Source data.** See the “Data availability” section.

**Competing interests.** The authors declare no competing interests.

---

**Extended Data Figure 1.**
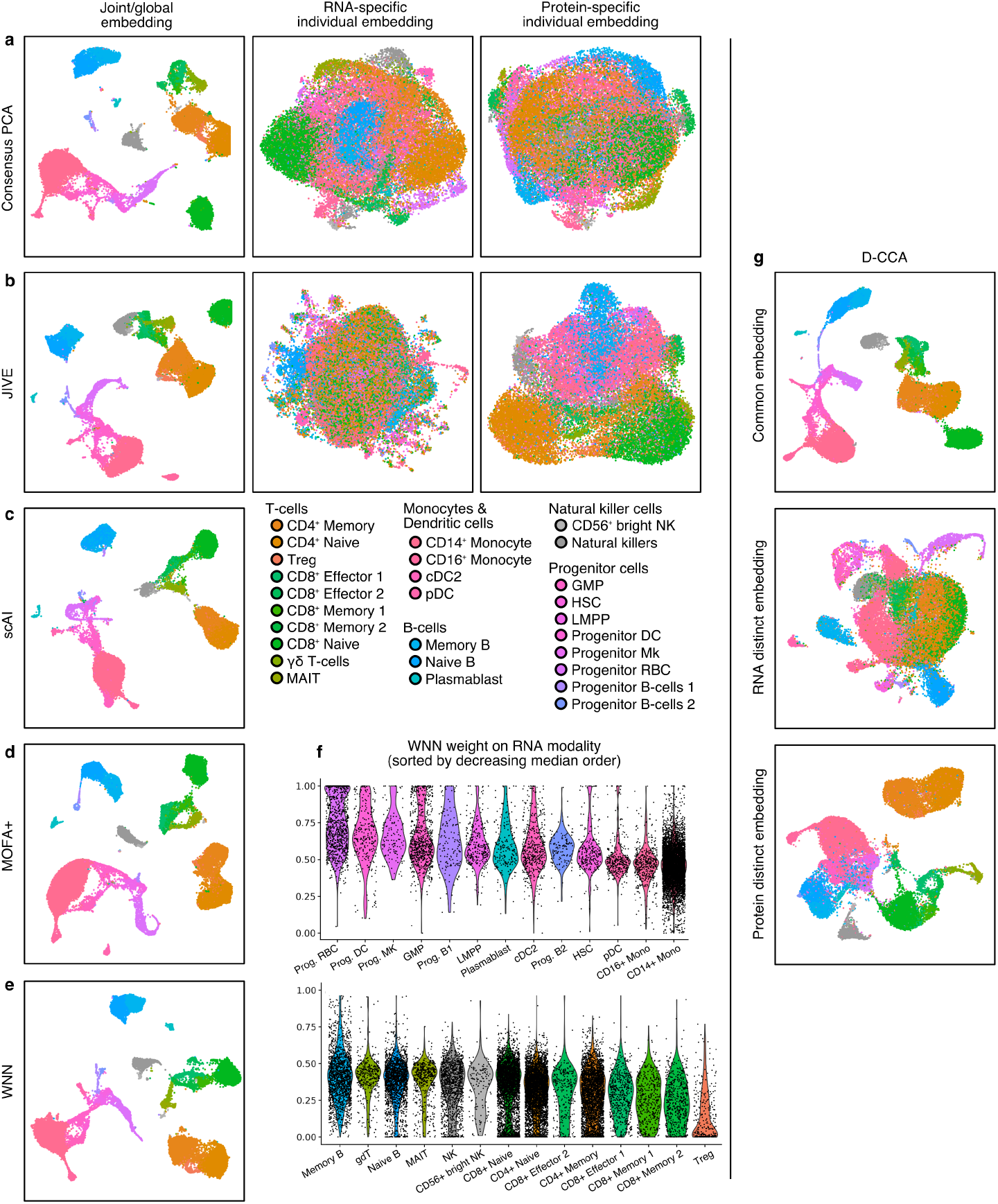
Embeddings of the human bone marrow CITE-seq data using various multiomic methods. **a**, Consensus PCA, showing the global embedding as well as RNA-specific and protein-specific residual embeddings. **b**, JIVE, showing the joint embedding as well as the RNA-specific and protein-specific individual embeddings. **c,d,e,**, scAI, MOFA+ and WNN, showing the global embedding. **f**, Violin plot of the modality-weight of cells computed by WNN, separated by cell-types, where a higher value denotes a cell having more informative neighbors in the RNA modality. Observe that younger/progenitor cells rely more on the RNA modality, while mature T-cells rely more on the protein modality. **g**, D-CCA, showing the common, RNA-distinct, protein-distinct embeddings. Observe that compared to Figure 1f, the common embedding for D-CCA separates CD4+ and CD8+ T-cells (which is a quality unique the protein modality), which can impact the downstream analyses.

**Extended Data Figure 2.**
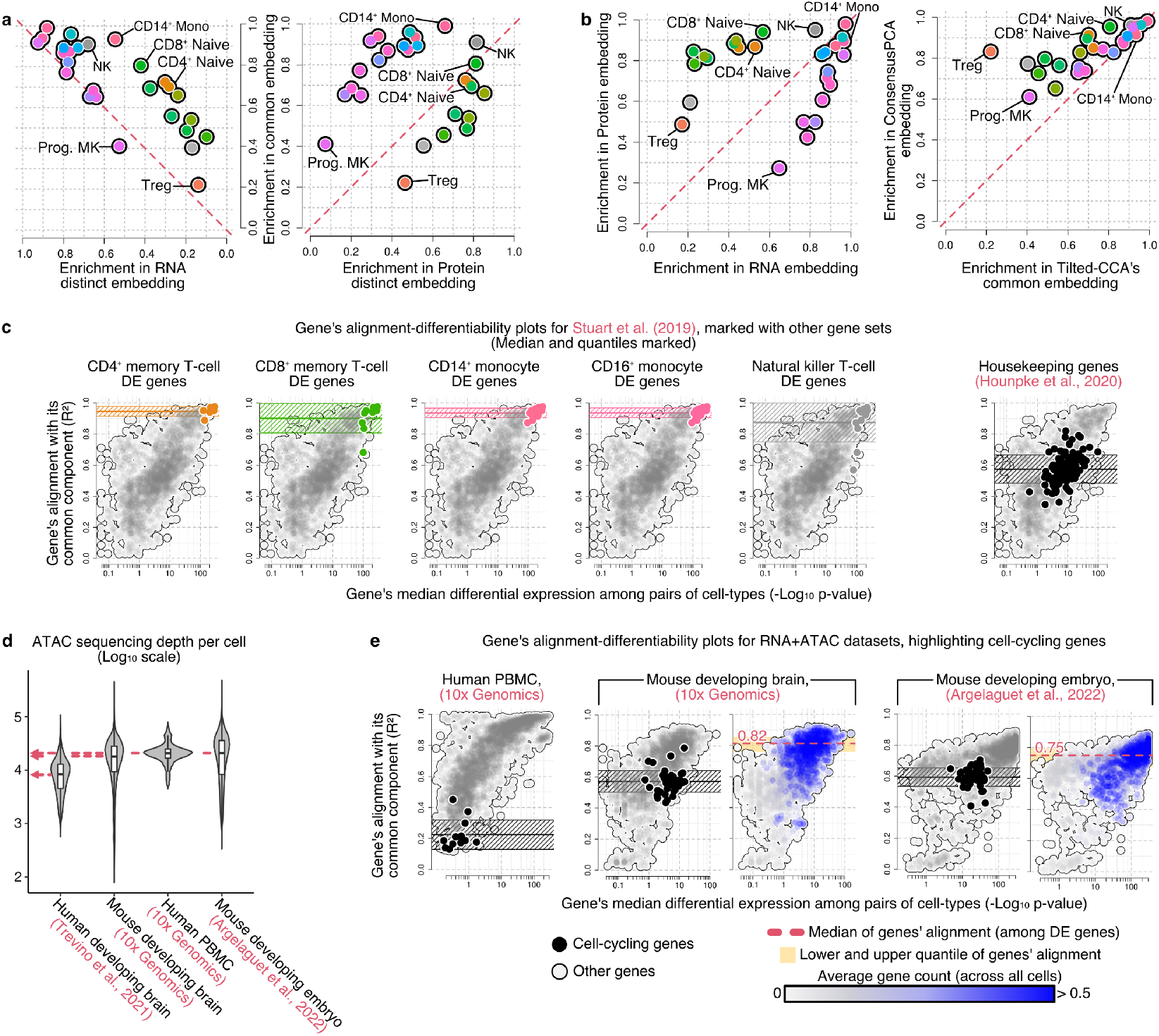
Additional plots illustrate the overlap/distinct information among cells or the alignment of genes to other modalities across different datasets. **a**, Enrichment of cell-types in the three embeddings for the human bone marrow CITE-seq data, showing either the enrichment of the common versus the RNA-distinct embedding (left) or common versus the protein-distinct embedding (right). **b**, Enrichment of cell-types in the RNA versus protein modality, analyzed separately (left), as well as in the Consensus PCA global embedding versus Tilted-CCA common embedding (right). Note in the former plot, the enrichment scores show that T-cell subtypes are more separable in the protein modality while the progenitor/myleoid subtypes are more separable in the RNA modality, and in the latter plot, Consensus PCA (which captures the “union” of information) yields an embedding that has a higher separation across all cell types than Tilted-CCA (which is desired, as Tilted-CCA captures the “intersection” of information). The *y* = *x* line is shown as a red dotted line as reference for each plot in (**a**, **b**). **c**, Additional alignment-differentiability plots (comparable to those in Figure 3(**c**)) for the human bone marrow CITE-seq data, highlight marker genes for different cell types or housekeeping genes.**d**, Sequencing depth for the ATAC modality across different multiomic datasets. **e**, Additional alignment-differentiability plots for RNA+ATAC datasets (comparable to those in Figure 3(**e**)), highlighting either the median alignment among genes associated with cell-cycling or among all DE genes.

**Extended Data Figure 3.**
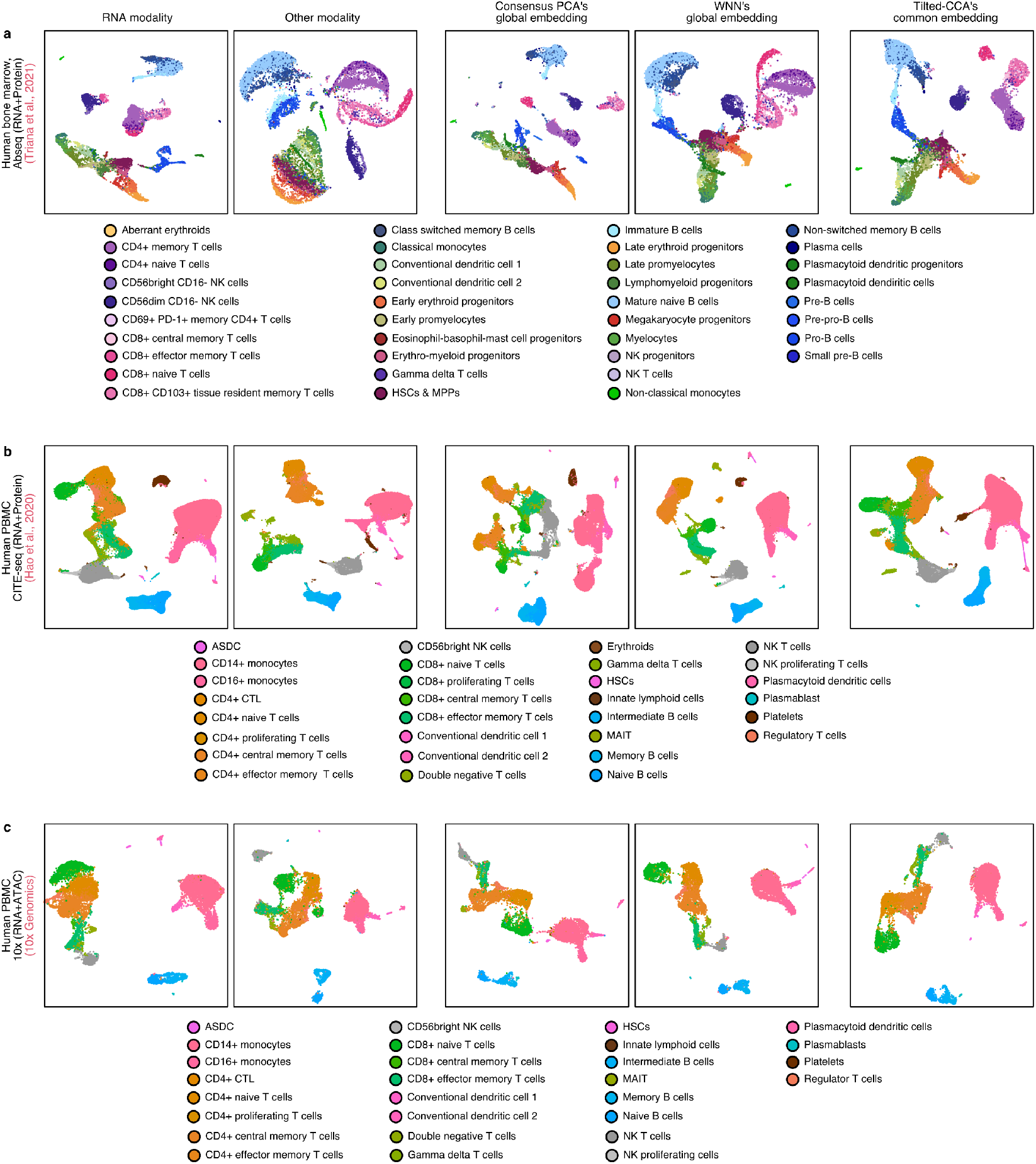
UMAPs of multiomic datasets of bone marrow and PBMC. **a**, Human bone marrow Abseq data jointly sequencing the RNA and protein modalities defined by 97 antibodies. **b**, Human PBMC CITE-seq data jointly sequencing the RNA and protein modalities defined by 224 antibodies. **c**, Human PBMC 10x data jointly sequencing the RNA and ATAC modalities. For each plot, the UMAP of the RNA modality alone, the other modality alone, Consensus PCA’s or WNN’s global embedding, and Tilted-CCA’s common embedding are shown, where cells are colored by the annotated cell-types for each dataset.

**Extended Data Figure 4.**
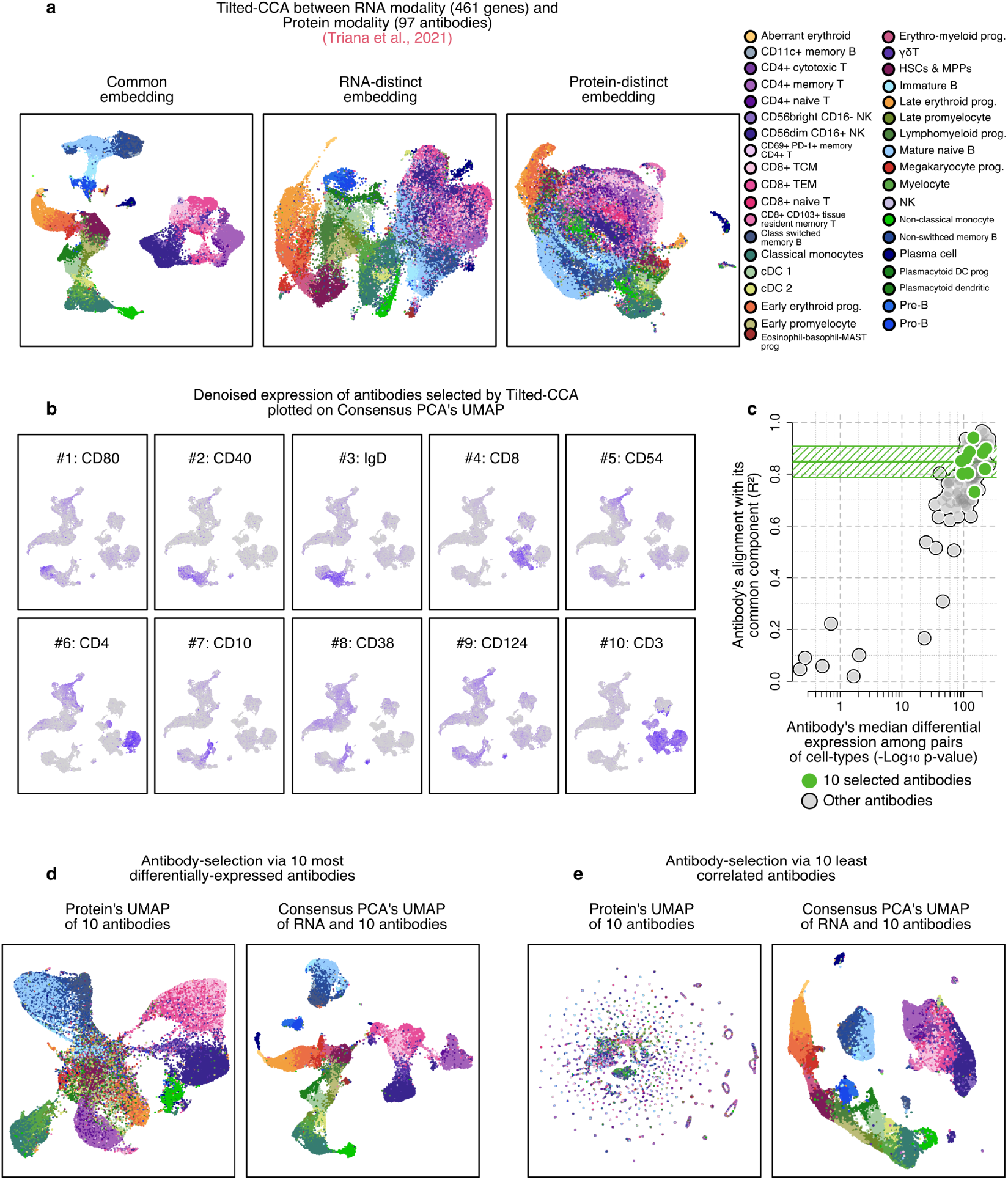
Additional plots to illustrate the antibody panel design case-study. **a**, UMAPs of the common, RNA-distinct and protein-distinct embeddings derived by Tilted-CCA when applied to the Abseq dataset jointly sequencing the RNA modality (defined by 461 genes) and protein modality (defined by 97 antibodies). We observe modality-specific axes of variation separating cell types in both distinct embeddings. Recall that Tilted-CCA ensures the axes of variation are orthogonal to one another. **b**, Feature plot showing the surface antibody marker expression for the 10 selected antibodies, overlaid on top of the Consensus PCA UMAP derived from the RNA modality and the 10 selected antibodies shown in Figure 4(**a**). **c**, Alignment-differentiability plot of the 97 antibodies, with the 10 selected antibodies marked. **d**, UMAP of the protein modality based on the 10 most differentially-expressed antibodies, as well as the Consensus PCA combining the axes of variation in the RNA modality and the 10 antibodies. Observe certain T-cell subtypes are not as separated in the Consensus PCA (compared to those in Figure 4(**a**)) since the 10 most differentially-expressed antibodies could contain axes of variation already found in the RNA modality. **d**, UMAP of the protein modality based on the 10 antibodies least correlated wih the RNA modality, as well as the Consensus PCA combining the axes of variation in the RNA modality and the 10 antibodies. Observe that these 10 antibodies do not axes of variation that help differentiate cell-types.

**Extended Data Figure 5.**
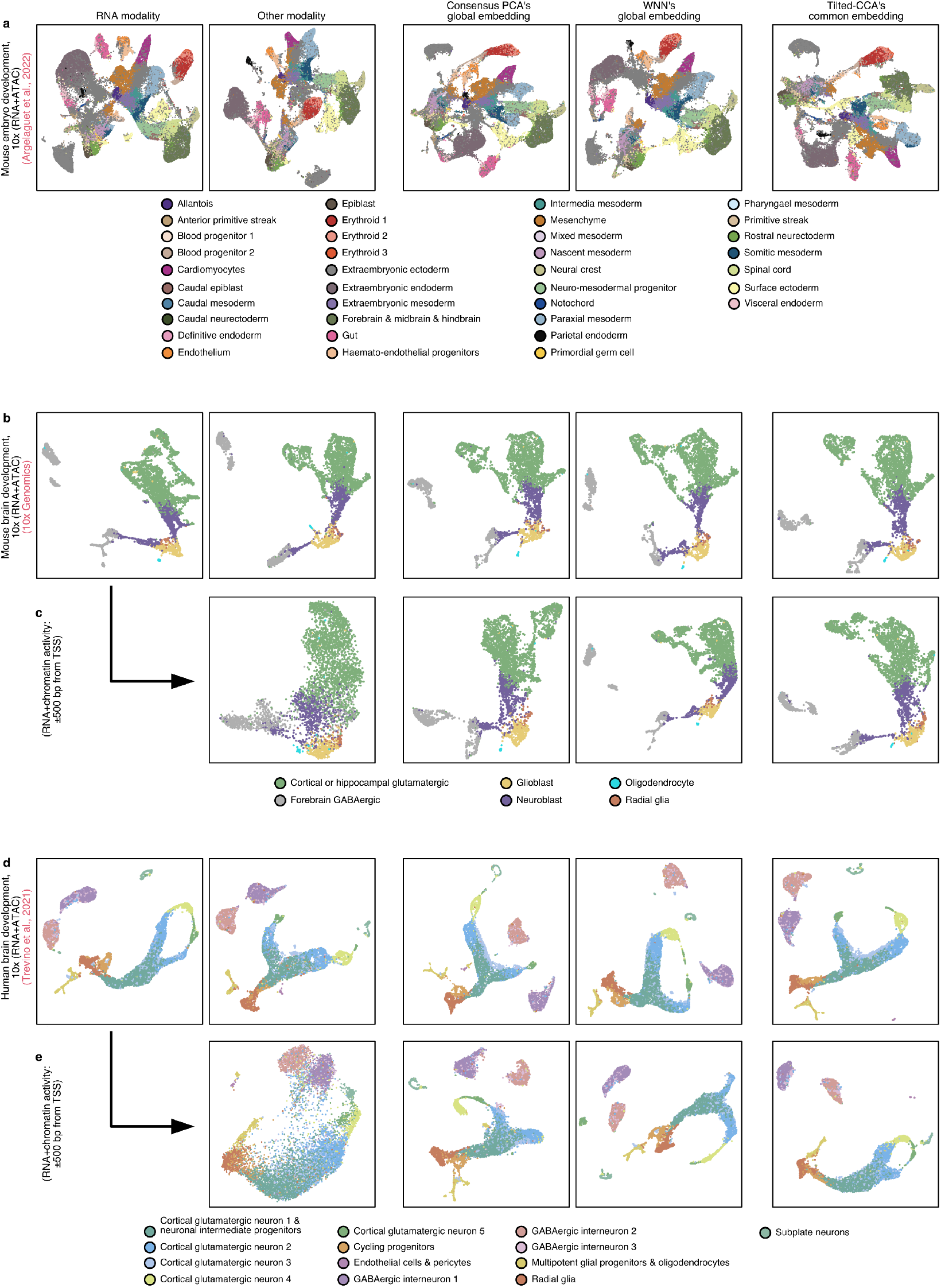
UMAPs of multiomic datasets sequencing RNA+ATAC. **a**, Mouse embryo development system, showing embeddings derived from the RNA and ATAC modalities. **b**, Mouse brain development system, showing embeddings derived from the RNA and ATAC modalities. **c**, Mouse brain development system, showing embeddings derived from the RNA and chromatin activity modalities. **d**, Human brain development system. **e**, Human brain development system, showing embeddings derived from the RNA and chromatin activity modalities. All three datasets are sequenced using the 10x multiome. For each plot, the UMAP of the RNA modality alone, the other modality alone, Consensus PCA’s or WNN’s global embedding, and Tilted-CCA’s common embedding are shown, where cells are colored by the annotated cell-types for each dataset.

**Extended Data Figure 6.**
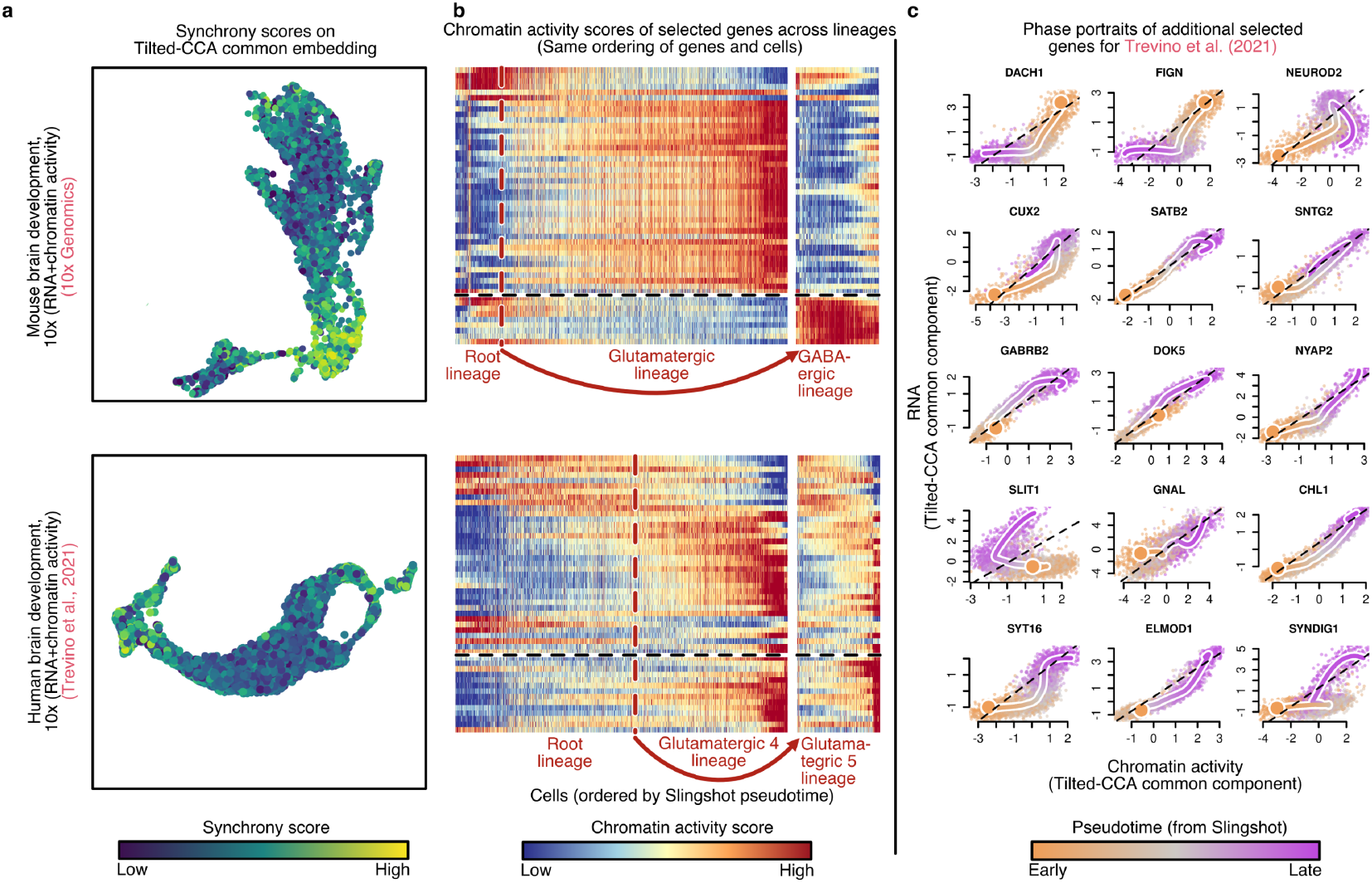
Additional plots supplementing findings of developmental qualities inferred using Tilted-CCA. **a**, The UMAPs of the common embedding between RNA and chromatin activity for the mouse brain development and human brain development system, where cells are colored based on their synchrony score. Observe that higher synchrony scores still imply cells in a terminal cell-state, but compared to the synchrony scores derived using RNA and ATAC shown in Figure 5(**d,f**), the signal is more feint. **b**, Heatmaps of the 50 selected development-associated genes and cell across pseudotime, organized in the same ordering as shown in Figure 5(**j,k**) but now showing each gene’s chromatin activity across various cells instead of each gene’s expression. **c**, Phase portraits of additional genes among the 50 selected development-associated genes for the human brain development system along the Glutamatergic 4 lineage, showing the relation between their gene expression and chromatin activity. These phase portraits supplement those in 6 to illustrate diversity of RNA-chromatin activity relations across development.

## 1 Brief description of other embedding methods for multiomic data

The following subsections describe the high-level ideas behind many of the dimension-reduction methods mentioned in the paper. These descriptions are not meant to reflect the specific nuances and details posed by the authors, but rather as a simplified sketch of each method to provide the reader a broad context on how each of the methods relate to one another. Please refer to the original text for each method for the exact details.

The theme is quite straightforward: loosely speaking, methods that aim to capture the “union of information” find a global embedding by maximizing its predictive power. On the other hand, methods like Tilted-CCA that aim to capture the “intersection of information” find a lower-dimensional embedding based on highly correlated signals in both modalities.

### 1.1 Consensus PCA

Consensus Principal Component Analysis (PCA), illustrated in Extended Data Figure 1a, comes in many forms historically in the literature, and we describe the specific variant used in this paper. Let 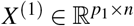 and 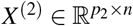 denote observed matrix for both modalities. Importantly, both *X*^(1)^ and *X*^(2)^ should be preprocessing in the following way: each feature is rescaled to have an empirical mean of 0 and an empirical standard deviation of 1. Then, each modality is rescaled by their operator (spectral) norm (i.e., the largest signular value). That is, for Modality *ℓ* ∈{1, 2},

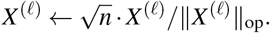

Finally, we denoise each modality (via PCA) by projecting *X*^(1)^ into a rank-*r*_1_ subspace and *X*^(2)^ into a rank-*r*_2_ subspace for pre-specified latent dimensionalities *r*_1_, *r*_2_.

Then, after this preprocessing, Consensus PCA applies PCA to the row-wise concatenation of *X*^(1)^ and *X*^(2)^. Specifically, it minimizes

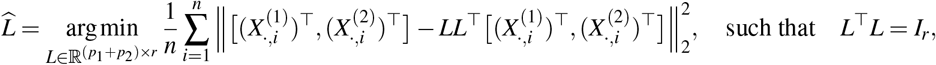

for a latent dimensionality *r* = min{*r*_1_, *r*_2_}, i.e., minimizing the reconstruction error. Observe that *LL*^⊤^ is a projection matrix that projects any *p*_1_ + *p*_2_ vector into a *r*-dimensional subspace. We can then treat

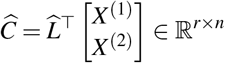

as the rank-*r* global embedding discovered by Consensus PCA, and the first *p*_1_ rows or last *p*_2_ rows of 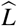 as Modality 1’s or Modality 2’s specific loading matrix respectively.

The modality-specific individual embeddings shown in Extended Data Figure 1a can then be computed by orthogonal component to 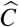. Specifically, let 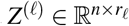 denote the leading principal components of *X*^*ℓ*^ for Modality *ℓ* ∈{1, 2}. Then, the modality-specific individual embedding for Modality *ℓ* is

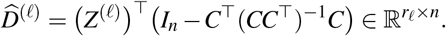

We deem that Consensus PCA as a method that captures the “union of information” since it finds a global subspace 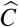 that minimizes the sum of reconstruction errors from each modality.

### 1.2 JIVE

Joint and Individual Variation Explained (JIVE)^10^, illustrated in Extended Data Figure 1b, is a linear matrix factorization method aimed to capture the “union of information” while providing an explicit decomposition of the common and distinct factors. Suppose 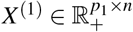 and 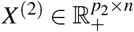 represent the preprocessed expression matrices of two different modalities. JIVE posits the model

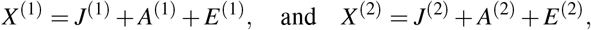

where 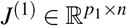 and 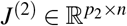 denote the joint components, satisfying

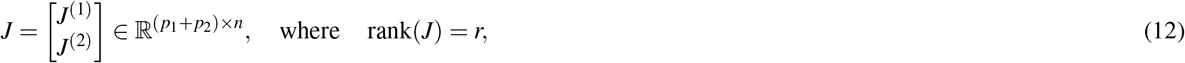

(i.e., is low-rank), and 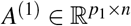 and 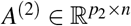 denote the modality-individual components where

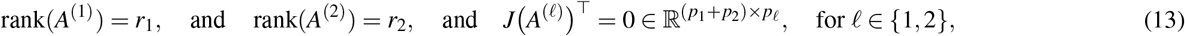

(i.e., the joint and modality-individual components are orthogonal) and 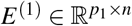 and 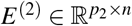 are matrices of zeromean independent noise. Here, *r, r*_1_, *r*_2_ are all pre-specified where *r* ≤ min{*r*_1_, *r*_2_}. Observe that the constraints in (12) and (13) can be satisfied if there exists matrix of basis vectors 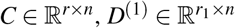 and 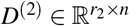 where

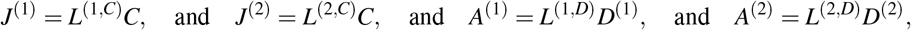

for various loading matrices 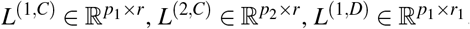, and 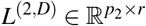, where importantly,

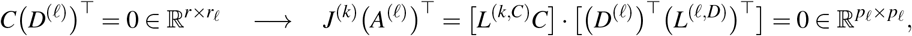

for any Modalities *k, ℓ* ∈ {1, 2}. Hence, this shows that JIVE strives to find lower-dimensional embeddings in a n-dimensional space where the basis vectors for the joint components are orthogonal to the basis vectors for either modality’s individual components.

JIVE estimates these components via an alternating-minimization scheme to solve

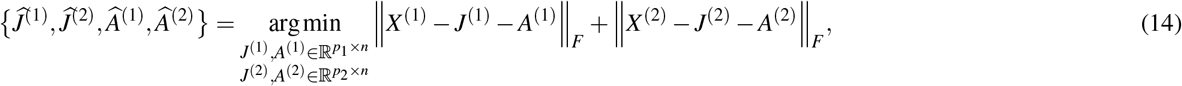

subject to

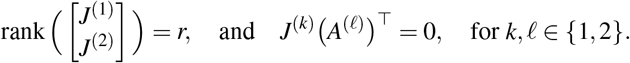

Similarly to Consensus PCA, it is important to appropriate scale *X*^(1)^ and *X*^(2)^ prior to using JIVE to ensure both modalities are “equally important.” In our work, for Modality *ℓ* ∈ {1, 2}, we first center all the features in *X*^(*ℓ*)^ to have an empirical mean of 0 and an empirical standard deviation of 1, and then normalize by

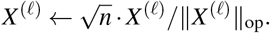

Since these low-rank matrices *J*^(1)^, *J*^(2)^, *A*^(1)^, *A*^(2)^ are estimated in (14) to minimize the prediction error (i.e., with additional rank constraints of the modality-specific components *A*^(1)^ and *A*^(2)^ not found in Consensus PCA), we deem the matrix *J* in JIVE as capturing the “union of information.”

### 1.3 scAI

Single-cell aggregation and integration(scAI)^13^, illustrated in Extended Data Figure 1c, is a linear non-negative matrix factorization method aimed to capture the “union of information” tailored specifically for multiomic data measuring the RNA and ATAC modalities. scAI first generates a random binary matrix *R* ∈ {0, 1}^*n* × *n*^ using a independent Bernoulli random variables with tuning paramter *s*, and strives to model the multimoic data as

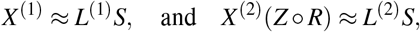

where 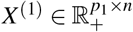 represents the observed raw counts of the RNA modality (i.e., not preprocessed or denoised), 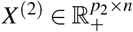 represents the observed raw counts of the ATAC modality, 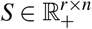 denotes the unknown non-negative global embedding of the *n* cells for a pre-specified latent dimensionality *r*, 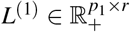 and 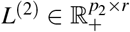 denote the unknown non-negative RNA-specific and ATAC-specific loading matrices, and *Z* is an unknown cell-cell similarity matrix that helps smooths the ATAC expression across cells to counteract the extreme sparsity of the ATAC modality.

To fit this model, scAI solves the following optimization problem,

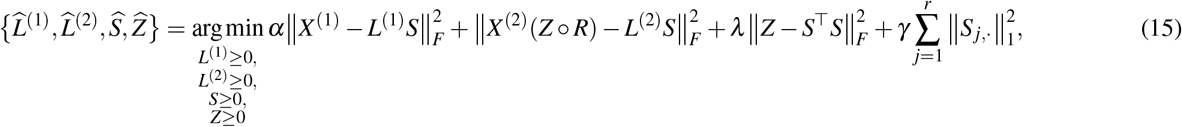

where *α, λ, γ* > 0 are tuning parameters and || ·||_*F*_ denotes the Forbenius norm. The terms 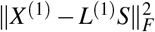 and 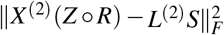 measure the quality of the matrix factorization, 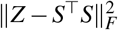 measures how well *Z* approximates the similarity among cells in *S*, and 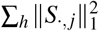 encourages sparsity in each latent factor. Additionally, the authors provide a procedure to interpret the loadings “shared” in both modalities based on *L*^(1)^ and *L*^(2)^, but as mentioned in the main text, we strive to design Tilted-CCA so the common signals can be inferred directly from the low-dimensional embedding.

When we use scAI on the bone marrow CITE-seq data^18^ in Extended Data Figure 1c, we instead optimize,

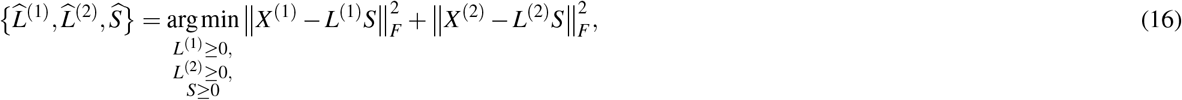

a simplification from (15) since the protein modality is more densely-sequenced compared to the ATAC modality.

We deem scAI as a method that captures the “union of information” since it finds a global subspace 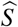 that minimizes the sum of reconstruction errors from each modality.

### 1.4 MOFA+

Multi-omics Factor Analysis (MOFA+)^12^, illustrated in Extended Data Figure 1d, is a linear matrix factorization method aimed to capture the “union of information” that is fit based on a Bayesian model where the generative model and priors are designed specifically for single-cell data. Suppose 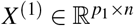 and 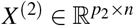 represent the expression matrices of two different modalities, preprocessed to be roughly Gaussian-distributed. Specifically, the generative model is

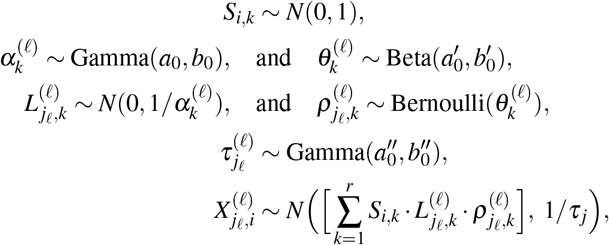

where *i* ∈ {1,…, *n*} denotes the index for cells, *k* ∈ {1,…, *r*} denotes the index for latent factors where *r* can be chosen in a data-driven fashion, *ℓ* ∈ {1, 2} denotes the index for the modality, and j_*ℓ*_ ∈ {1,…, *p_ℓ_*} is the index for features in the *ℓ*th modality, and *a*_0_, *b*_0_, 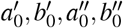 all represent hyperparameters chosen beforehand. Here, *S_i,k_* denotes the cell *i*’s global embedding in latent dimension *k*, and 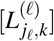 and 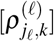 act as the factor matrix specific for Modality *ℓ*, where the priors are chosen to mimic a spike-and-slab prior (to induce sparsity in the factors) while be amendable for tractable Bayesian inference.

We deem MOFA+ as a method that captures the “union of information” since it models both modalities as stemming from a global embedding *S* where modality-specific factors 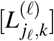 and 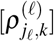 “pick out” the relevant factors to reconstruct each modality, and *S* is optimizes to minimize the prediction errors.

### 1.5 WNN

Weighted Nearest-Neighbor (WNN)^11^, illustrated in Extended Data Figure 1e, is a nonlinear embedding method for multiomic data aimed to capture the “union of information” based on nearest-neighbor graphs. Let 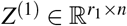 and 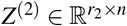 denote leading principal components of *X*^(1)^ and *X*^(2)^ (after being appropriately preprocessed). WNN first constructs *G*^(1)^, *G*^(2)^ ∈{0, 1}^*n* × *n*^, the two nearest-neighbor graphs from each modality. Then, WNN compares how well each cell’s expression in each modality can be predicted by the average expression of that cell’s neighbors from either graph. Specifically, for cell *i* ∈ {1,…, *n*} let 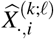 denote the average expression of cell *i* in Modality *k* using the nearest neighbors of cell *i* dictated by *G^ℓ^*, for *k*, *ℓ* ∈ {1, 2}. Then, let

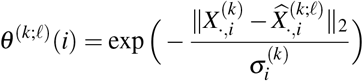

denote how well the nearest neighbors from Modality *ℓ* predict the expression of cell *i* in Modality *k*, using a Gaussian kernel with an appropriately chosen bandwidth 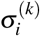. Then, based on how *θ*^(*k*;*k*)^ (*i*) compares to *θ*^(*k*;*ℓ*)^ (*i*) for *k, ℓ* ∈ {1, 2} (i.e., is a cell’s expression better predicted using the nearest-neighbors in its native modality than the nearest-neighbors in other modality), WNN computes 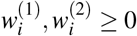 where 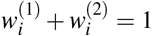, two scalars that weights the relative importance of each modality for understanding cell *i*’s expression. Finally, WNN outputs the weighted graph 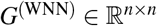 where, for any pairs of cells *i, j* ∈ {1,…, *n*}

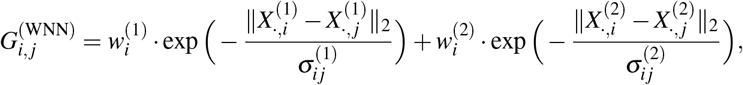

for appropriately chosen bandwidths 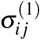 and 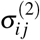. Then, any embedding method for graphs can be applied on *G*^(WNN)^ to extract a relevant low-dimensional embedding.

We deem WNN as a method that captures the “union of information” since by design, if Modality 1 has more informative neighbors than Modality 2 for cell *i*, then *G*^(WNN)^ would rely on Modality 1 more than Modality 2 to determine the appropriate similarity-scores between cell *i* and all other cells.

### 1.6 D-CCA

Decomposition CCA (D-CCA)^39^, illustrated in Extended Data Figure 1g, is a predecessor of Tilted-CCA and is exactly Tilted-CCA when *τ_j_* = 0.5 for all latent dimensional *j* ∈ {1,…, *r*}. Specifically, after computing the canonical scores *Z*^(1)^, 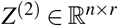, D-CCA decomposes each pair of canonical score vectors *j* ∈ {1,…, *r*} by solving

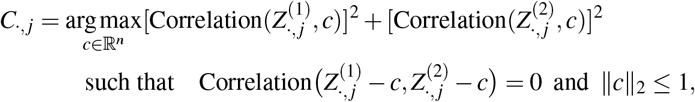

where Correlation(·, ·) denotes the Pearson correlation between two vectors. The above optimization is solved by setting *τ_j_* = 0.5 in our framework, where the common vector *C*._,*j*_ perfectly bisects 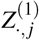 and 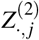 on the 2-dimensional hyperplane where both vectors lay ontop of (see Figure 1H). This was showed by the authors of D-CCA, where the closed-form solution to the above optimization

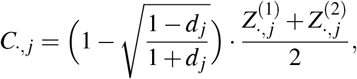

is where *d_j_* ∈ [0, 1] denotes the *j*th canonical correlation between the two modalities. Recalling that 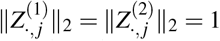 since they are canonical score vectors, observe that this also shows for the specific choice of *τ_j_* = 0.5, the length of the common vector *C*._,*j*_ (i.e., ||*C*.,*j*||_2_) is (non-linearly) proportional to the canonical correlation *d_j_*. Observe that in contrast to the aforementioned methods that capture the “union of information,” here, the embedding *C* is designed not to minimize the prediction error but rather to extract correlated signals from both modalities.

We are motivated to develop Tilted-CCA because while D-CCA provides a sensible way to construct *C*, there is no inherit geometric property among C that captures the “intersection of information” intuition we seek for our biological applications. Hence, this is why we relax the choice of *τ_j_* = 0.5 and instead optimize over *τ_j_* ∈ [0, 1] such that the cells in *C* have geometric relations that reflect the target common manifold *G*. This way, the distinct vectors 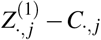 and 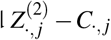 are still orthogonal in each latent dimension (i.e. preserving the interpretation of axes of variation unique to a modality) while the geometric relations in *C* are themselves informative.

## 2 Inferring the common/distinct axes of information from embeddings that capture the “union of information”

Note that for certain matrix factorization methods for multiomic data, it is possible to infer which axes of variation in the joint embedding are “shared” between both modalities or “unique” to a particular modality by investigating the features’ loadings across latent factors. This is best exemplified by scAI, since its non-negative joint matrix factorization enables interpretable loadings *L*^(1)^ and *L*^(2)^. Consider the scAI embedding of the human bone marrow CITE-seq data shown in Extended Data Figure 1c. We plot the mean loading for each of the 30 latent factors across all the features (i.e., genes in the RNA modality and antibodies in the protein modality) (Supplementary Fig. 1a). Both modalities have relatively the same loading for Factor 4, suggesting this factor denotes an axes of variation shared between both modalities. When investigating the scores for this factor across all the cells, we see that this latent factor represents an axes of variation that separates the B-cells (blue) from other cell types (Supplementary Fig. 1b). On other hand, the loadings for Factor 19 in the protein modality is dramatically higher than those in the RNA modality in the RNA modality, which suggests this factor denotes an axes of variation unique to the protein modality. The score along this factor separate the CD4+ T-cells (orange) from the other cells (Supplementary Fig. 1c). Lastly, the loadings for Factor 20 in the RNA modality is dramatically higher than those in the RNA modality in the protein modality, which suggests this factor denotes an axes of variation unique to the RNA modality. The score along this factor separate the myeloid cells (purple) from the other cells (Supplementary Fig. 1d). All these findings for B-cells, CD4+ T-cells and myeloid cells match our enrichment analysis provided in Figure 2a. The authors of scAI also suggest similar diagnostics to infer the common/distinct axes of variation between both modalities.

**Supplementary Figure 1.**
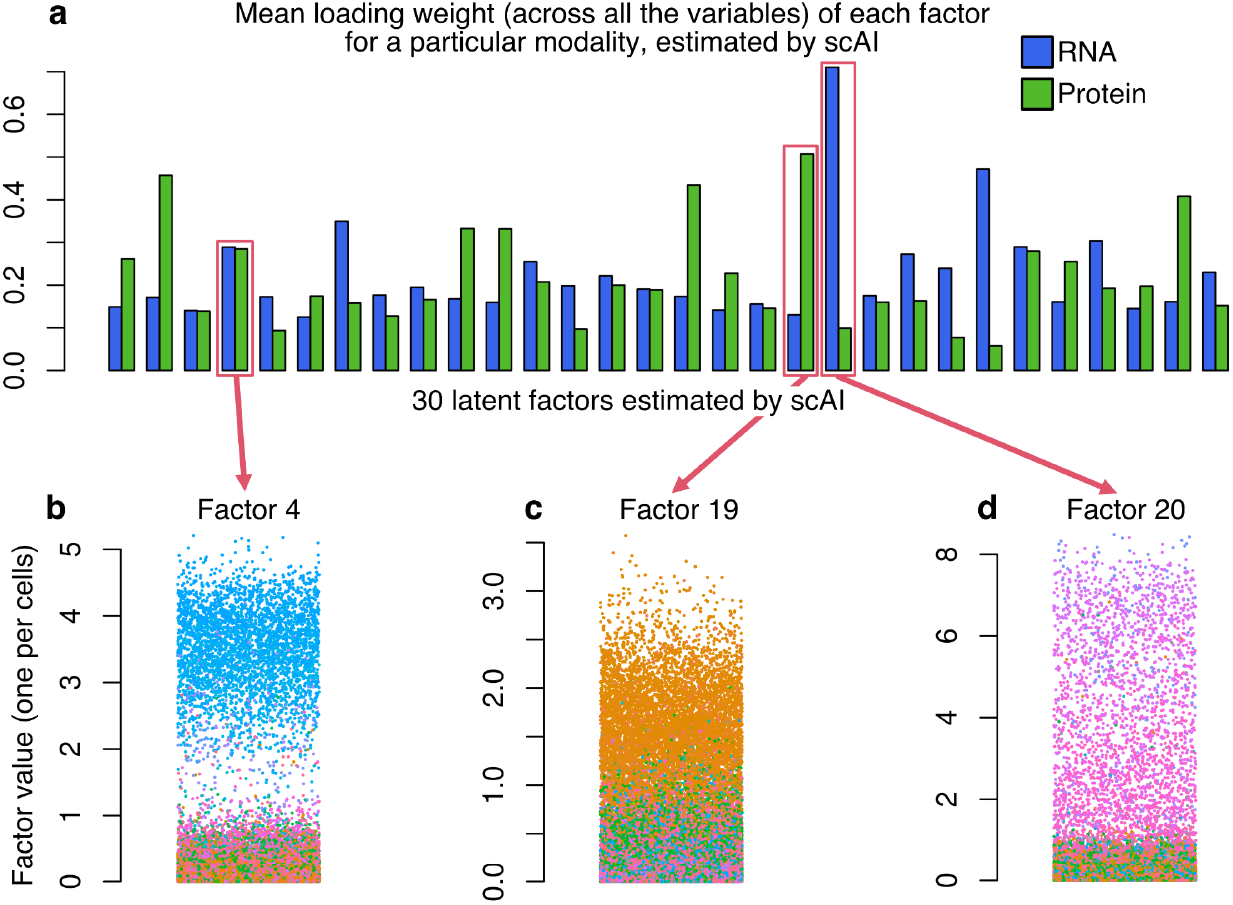
Illustration of inferring the common/distinct information using a matrix factorization method that captures the “union” of information. **a**, The mean loading across all the features in either the RNA (blue) or protein (green) modalities for all 30 factors, where we focus on three factors in particular (bottom). **b**, The weights across the cells in Factor 4 illustrates a separation of B-cells from all other cells, and both modalities weigh this factor equally, suggesting a “common” axes of variation. **c,d**, The weights across the cells in Factor 19 or 20 illustrate a separation of CD4+ T-cells or certain myeloid cells from all other cells respectively, and the mean weight of these factors suggest that Factor 19 is an axes of variation “unique” to the protein modality while Factor 20 is an axes of variation “unique” to the RNA modality. Refer to Figure 1(**a**) for the correspondence between cell type and color.

However, despite matrix factorization methods such as scAI focusing on the “union” of information being able to deduce the “intersection” of information, such procedures are hindered by two aspects: 1) interpreting the common/distinct axes of variation requires a careful analysis of the loadings *L*^(1)^ and *L*^(2)^ in conjunction with the joint embedding S, whereas Tilted-CCA directly separates the common/distinct axes of variation through the embeddings *C, D*^(1)^ and *D*^(2)^ (which can each be visualized and studied in isolation), and 2) the aforementioned analyses of scAI’s joint embedding *S* does not explicitly separate and quantify the amount of common/distinct information between the two modalities. Such quantification would necessarily require some decomposition similar to Tilted-CCA’s decomposition described in Figure 1b, and we believe that Tilted-CCA’s framework enables the most direct and interpretable procedure to achieve this decomposition.

## 3 Mathematical details on constructing the target common manifold

### 3.1 Scenario with no global clustering structure

Suppose we are given the two nearest-neighbor graphs from each modality, *G*^(1)^, *G*^(2)^ ∈ {0, 1 }^*n × n*^. Then, for cell *i* ∈ {1,…, *n*}, let *A, B* ∈ {1,…, *n*}\{*i*} denote the set of edges involving cell *i* in either *G*^(1)^ and *G*^(2)^ respectively. Our goal is to construct a set *C* ⊆ *A* ∪ *B* that is the “midpoint” between the sets *A* and *B*, specifically one that satisfies

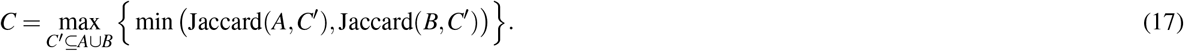

This is qualitatively similar to defining a midpoint *z* of two scalars *x, y* as *z* = max_*z*_′{min(|*x* – *z*2|, |*y* – *z*′|)}. The set *C* is not unique, but the cardinality of its components is (given regularity conditions). Specifically, let *s* =|*A* ⋂ *B*|, *d*_1_ = |*A*\*B*| and *d*_2_ = |*B*\*A*|, denoting the number of edges shared between *G*^(1)^ and *G*^(2)^, unique to *G*^(1)^, or unique to *G*^(2)^ respectively. We make two observations: first, optimizing (17) reduces to optimizing

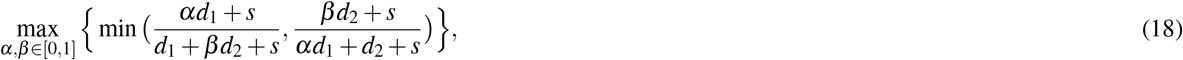

and then constructing *C* as all *s* edges in *A* ⋂ *B*, *α*% of the *d*_1_ edges in *A*\*B*, and *β*% of the *d*_1_ edges in *B*\*A*. Second, if *d*_1_ < *d*_2_ without loss of generality, then *α* = 1 and *β* < 1. Solving (18) amounts to solving a quadratic equation for *β* (since *α* = 1 without loss of generality). This quadratic equation is derived from (18), since the optimum must satisfy

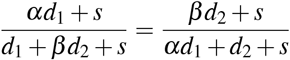

where *α* = 1.

### 3.2 Scenario with global clustering structure

Suppose we are given the two nearest-neighbor graphs from each modality, *G*^(1)^, *G*^(2)^ ∈ {0, 1}^*n ×n*^, two “hard” clustering vectors *c*^(1)^ ∈ {1,…, *m*_1_}^*n*^ and *c*^(2)^ ∈{1,…, *m*_2_}^*n*^, and a desired number of edges *k* for each cell. Then, for cell *i*∈{1,…, *n*}, let *A*, *B* ∈{1,…, *n*}\{*i*} denote the set of edges involving cell *i* in either *G*^(1)^ and *G*^(2)^ respectively. Additionally, for cell *i*, let *P*_1_: {1,…, *n*} → [0,1]^*m*_1_^ and *P*_2_: {1,…, *n*} →[0, 1]^*m*_2_^ be two functions that take in a set of indicies (i.e., neighboring cells to cell *i*) and outputs a multinomial vector that dictates the percentage of cells connected to cell *i* that belong to any of the *m*_1_ clusters in *G*^(1)^ or to any of the *m*_2_ clusters in *G*^(2)^ respectively (i.e., a non-negative vector that sums to 1).

Our goal is to construct a set *C* ⊆ *A* ∪ *B* that is the “midpoint” between the sets *A* and *B*, specifically one that satisfies

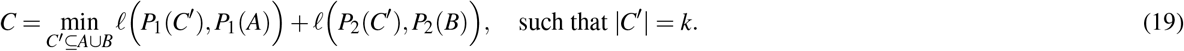

where *ℓ*(·, ·) is the squared *ℓ*_2_ distance between two multinomial vectors of the same dimensionality, i.e., for *x* = (*x*_1_,…,*x_m_*) ∈ [0,1]^*m*^ and *y* = (*y*_1_,…, *y_m_*) ∈[0,1]^*m*^ (both summing to 1),

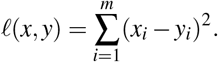

Similar to setting with no global clustering structure, the set *C* is not unique, but the cardinality of its components is (given regularity conditions).

To warm up to the solution, consider the following simplifying scenario that illustrates the nuisance of constructing *C*. Suppose *m*_1_ = 2, *m*_2_ = 3, and for a particular cell *i*, |*A*| = 30 where all 30 edges connect cell *i* with the first of the *m*_1_ = 2 clusters in Modality 1 (i.e., *P*_1_(*A*) = (1,0), meaning cell *i* is perfectly separated into one cluster in Modality 1), while |*B*| = 30 where 10 edges connect cell *i* with each of the *m*_2_ = 3 clusters (i.e., *P*_2_(*B*) = (1/3, 1/3, 1/3), meaning cell *i* lies between the three clusters in Modality 2). Additionally, suppose |*A* ⋂ *B*| = 0, but all 30 cells connected to cell *i* in Modality 2 are also in first of the *m*_1_ clusters in Modality 1, and all 30 cells connected to cell *i* in Modality 1 are also equally split among the *m*_2_ clusters in Modality 2. Then, consider the naive construction of *C* = *A* ∪ B, resulting in connecting cell *i* to 60 other cells. Under this construction, while *P*_1_(*C*) = (1,0), we see that *P*_2_(*C*) = (2/3, 1/6, 1/6). This is potentially concerning, as if we used this set *C* for cell *i* when constructing the target common graph *G*, cell *i* is now vastly more connected with one of the three clusters (from the perspective of Modality 2). This could have harmful consequences when *G* should ideally capture the “common” information between the two modalities. Arguably, in this scenario, if Modality 2 perceives cell *i* as “in between” three clusters, the target graph *G* should reflect this as well.

We describe the quadratic optimization program to solve (19) for a particular example when *m*_1_ = 2 and *m*_2_ = 3 for simplicity, from which the generalized procedure to solve (19) could be easily inferred. Let *P*_1_(*A*) = (*α*_1_, *α*_2_) and *P*_2_(*B*) = (*β*_1_, *β*_2_, *β*_3_), and let *n_i_j* denote the number of cells in *A* ∪ *B* (i.e., cells neighboring cell *i* in either *G*^(1)^ or *G*^(2)^) that are simultaneously in the *i*th cluster in Modality 1 and the *j*th cluster in Modality 2. We can visualize this scenario with the following confusion table,

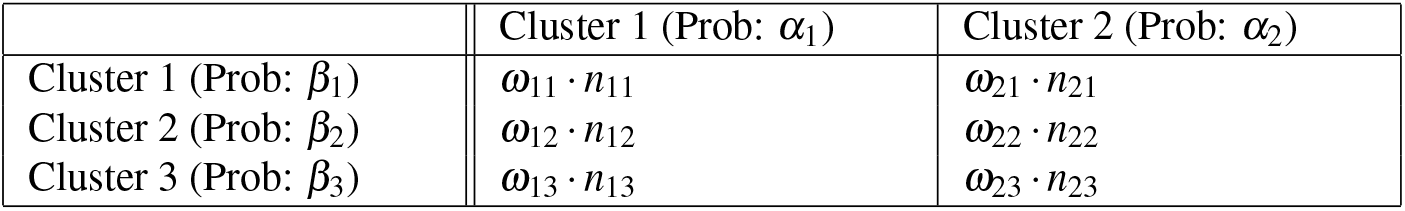

where the goal is to solve for scalars *ω*_11_, *ω*_12_,…, *ω*_23_ ∈ [0,1] minimizing a certain objective function (to be described) under the constraint

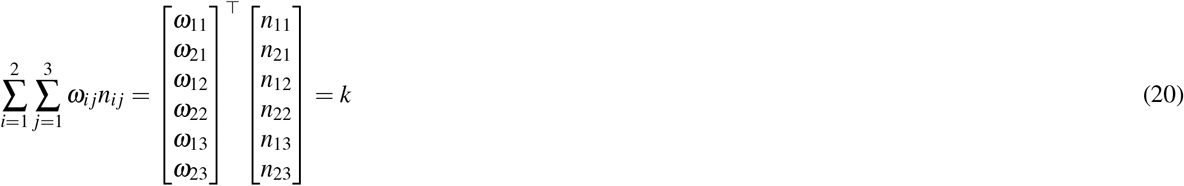

Once we have solved for *ω*_11_, *ω*_12_,…, *ω*_23_, the set *C* would be constructed by sampling *ω*_11_% of the *n*_11_ cells in Cluster 1 in Modality 1 and Cluster 1 in Modality 2, *ω*_21_% of the *n*_21_ cells in Cluster 2 in Modality 1 and Cluster 1 in Modality 2, and so on.

All that remains is to describe the objective function to the quadratic program. In the full generality, there are *m*_1_ + *m*_2_ major terms to consider, of which we will describe how to construct one of these *m*_1_ + *m*_2_ = 2 + 3 = 5 terms in our example. Observe that the objective function in (19) involves one term that computes if the proportion of elements in *C* that lie in Cluster 1 in Modality 1 (as dictated by *c*^(1)^) is close to *α*_1_. In our example, this is computed by

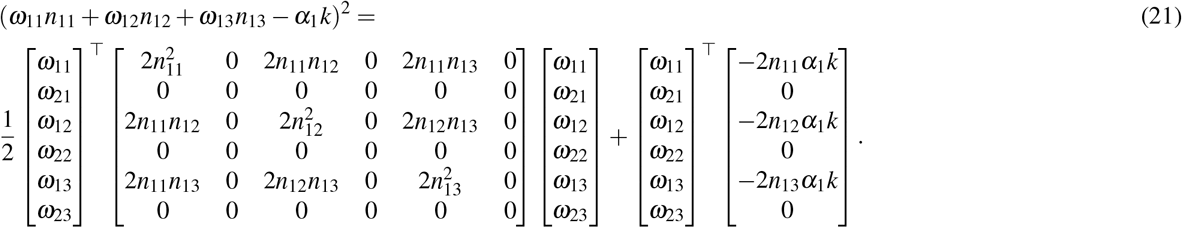

Observe that all *m*_1_ + *m*_2_ terms (of which one is shown in (21)) and the constraint in (20) are in the form of a quadratic optimization program. After formulating this, we solve this optimization program via `quadprog::solve.QP`, which informs us of the scalars *ω*_11_, *ω*_12_,…, *ω*_23_, which in turn allows us to construct the set *C* for cell *i* as described above.

## 4 Additional qualifications of RNA+protein data via alignment-differentiability plots

We use alignment-differentiability plots to investigate how increasing the antibody panel size impacts the separation of immune cell types using three RNA+protein multiome datasets: an Abseq dataset of bone marrow with 97 antibody markers^20^ (the “Triana” dataset), the aforementioned CITE-seq dataset of bone marrow with 25 antibody markers^18^ (the “Stuart” dataset) and a CITE-seq dataset of PBMC with 224 antibody markers^11^ (the “Hao” dataset). The alignment-differentiability plot of the first two datasets are shown in Supplementary Figure 2a, while that for the Hao dataset was shown in Figure 3e. We summarize the global RNA feature alignment to the common space by the median alignment score among genes with differentiability score of 10 or more (red dashed line). We perform two different analyses to compare these three datasets.

First, since these three datasets contain different sets of antibody markers, we also analyze the relation between the transcriptome and the same set of 25 antibodies from the Stuart dataset for all three datasets using Tilted-CCA, when we apply Tilted-CCA on the same 25 antibodies from the Stuart dataset for all three datasets. In this scenario, we observe that the PBMC system has a higher mean alignment than either datasets of the bone marrow system (0.77 compared to 0.66 or 0.72, Supplementary Fig. 2a,b). This is expected since the chosen 25 antibody markers are geared to separate differentiated immune cells, which are more prevalent in PBMC than in bone marrow^18^.

Second, when we apply Tilted-CCA to the Triana and Hao datasets using the full antibody panel sizes of 97 and 224 respectively instead of the limited panel size of 25, we observe that for both the Triana and Hao datasets, the median alignment increases when the analysis on the aforementioned set of 25 antibodies were instead applied to full panel sizes of 97 or 224 antibodies (from 0.66 to 0.83 and from 0.77 to 0.83 respectively) (Fig. 3e and Supplementary Figure 2b). This demonstrates that the majority of the additional differentially-expressed antibodies in both datasets reinforces existing expression patterns already present in the transcriptome. Additionally, the alignment-differentiability plots of the antibodies offer complementary evidence of this phenomenon – the 25 antibodies across all three datasets are highly informative of cell type, while they differ in degrees of alignment with the RNA modality (Supplementary Fig. 2c)

**Supplementary Figure 2.**
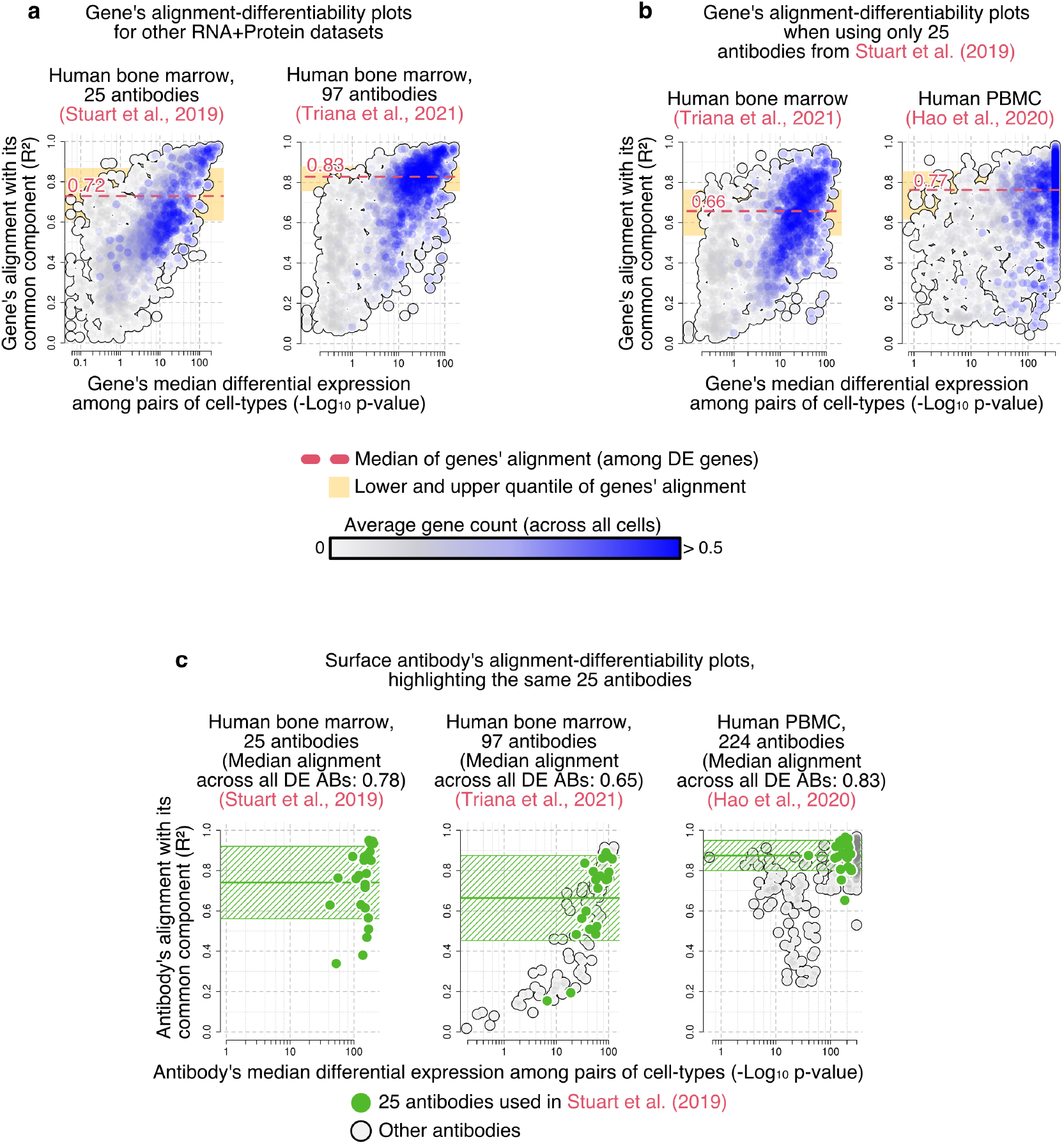
Alignment-differentiability plots for RNA+protein datasets that illustrate the impact of having additional antibodies in the panel. **a**, Additional alignment-differentiability plots for two datasets: human bone marrow sequenced via CITE-seq on 25 antibodies (i.e., the “Stuart” dataset) or Abseq on 97 antibodies (i.e., the “Triana” dataset). These complement the alignment-differentiability plot of the PBMC CITE-seq data of 224 antibodies in Figure 3(**e**) (i.e., the “Hao” dataset). **b**, Alignment-differentiability plot of the Triana or Hao dataset when applying Tilted-CCA on the RNA modality and the protein modality represented using only the 25 antibodies used in the Stuart dataset. Observe that the median alignment among DE genes in both datasets (red dotted lines) is lower compared the alignment derived from using each data’s full set of antibodies. **c**, Alignment-differentiability plot for all three dataset’s antibodies, where the same set of 25 antibodies from the Stuart dataset are marked in each plot.

